# Identifying membrane-bound transcriptional regulatory proteins from rare but evolutionarily conserved domain combinations

**DOI:** 10.64898/2025.12.20.695554

**Authors:** Muyoung Lee, Yeejin Jang, Qingqing Guo, Jonghwan Kim, Edward M. Marcotte

## Abstract

Transcriptional regulatory proteins, including transcription factors (TFs) and chromatin modifiers, must act inside cell nuclei, but membrane-bound transcription factors (MBTFs) are first anchored in membranes before nuclear translocation. Known MBTFs are vital for processes from myelin expression (MYRF) to cholesterol homeostasis (SREBP), yet their overall diversity remains uncharted. We hypothesized that additional membrane-bound transcriptional regulators (MBTRs) might exist, so we developed a bioinformatics screen to prioritize membrane proteins that are likely to regulate transcription. Our approach leverages domain composition by positing that surprising domain combinations suggest novel biological functions. We searched for rare but evolutionarily conserved pairings of transmembrane domains with domains likely involved in transcriptional regulation. Our method rediscovered known MBTFs and membrane-bound histone kinases, and identified novel MBTR candidates, including transmembrane histone N-acetyltransferases, a putative new subclass of MBTFs that share MYRF’s DNA-binding domain, and the prolactin regulatory element-binding protein PREB (SEC12). Assays using recombinant PREB demonstrated that the transmembrane domain is the determinant of PREB subcellular localization, governing its distribution between the membrane and the nucleus in mouse embryonic stem cells. These findings underscore the utility of our method and provide a framework for investigating MBTRs that likely facilitate the integration of extracellular signals with transcriptional responses.

**Graphical Abstract:** 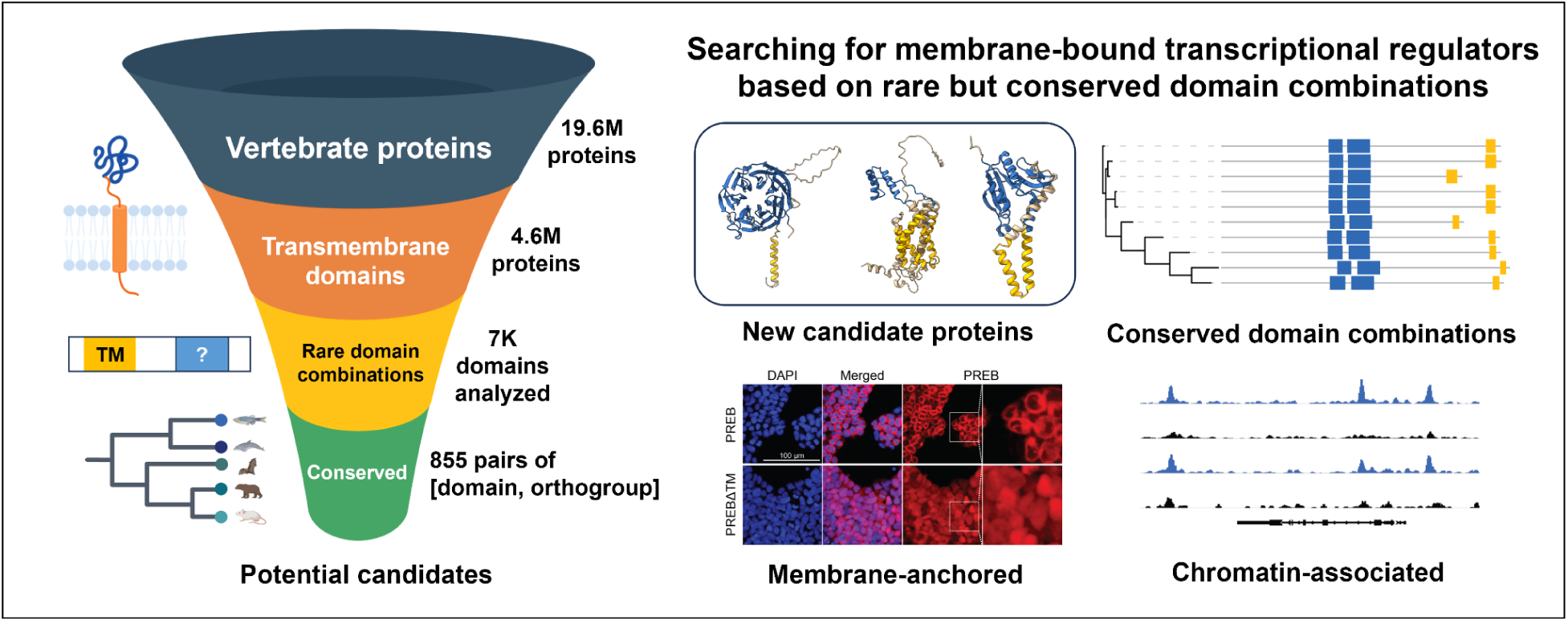

## Introduction

Transcription factors (TFs) regulate transcription by binding to DNA in the nucleus (1), so many TFs are imported into the nucleus from the cytosol. However, some TFs are anchored to membranes by transmembrane domains before their nuclear translocation. These rare TFs, known as membrane-bound transcription factors (MBTFs), play crucial biological functions. For example, SREBP (Sterol regulatory element-binding protein) regulates lipid metabolism and homeostasis (2), MYRF (Myelin regulatory factor) regulates myelin production by glial cells (3), and ATF6 (Activating transcription factor 6) reacts to the accumulation of misfolded proteins in cells (4). For an MBTF to function as a TF, it must be released from the anchored membrane and transported to the nucleus. Multiple mechanisms are known for this proteolytic release from the transmembrane domains (3, 5, 6).

Efforts to systematically screen for MBTFs have been ongoing, with the first computational prediction of MBTFs published in 2001 (7). This study identified proteins with selected DNA-binding domains from the Pfam database (8) and transmembrane domains using a consensus of multiple prediction tools. From 20 organisms and one virus, they identified 76 proteins, including six human proteins such as SREBP1, SREBP2, and DMRT2. However, they excluded ATF6 because only two of the four programs predicted its transmembrane domain, and proteins lacking the selected DNA-binding domains were also filtered out. Yao et al. identified 1,089 MBTFs from a total of 25,850 TFs in 14 plant species, including 64 MBTFs from *Arabidopsis thaliana* (9). They predicted the presence of transmembrane domains in transcription factors from PlantTFDB v4.0 (10). While both studies predicted the existence of transmembrane domains in proteins with selected DNA-binding domains or TFs from a database, the actual number of MBTFs remains largely unknown. Furthermore, it is uncertain how common these types of transcriptional regulation are, including those involving membrane-bound proteins that bind DNA (e.g., MBTFs), modify chromatin, or perform previously undiscovered functions.

In this study, we developed a protein-domain-based search method to identify membrane-bound transcriptional regulators (MBTRs), including both MBTFs and membrane-bound chromatin modifiers, without imposing any specific precondition on the target domain characteristics. Many proteins consist of structurally and functionally independent domains that play key roles in determining a protein’s overall function. We hypothesized that by identifying proteins with domains that are prevalent in non-membrane proteins yet scarce in membrane proteins, we might uncover new MBTRs. Given the importance of known MBTFs in vertebrate embryonic development and health, we focused exclusively on vertebrate proteins, aiming to identify new conserved human and mouse MBTRs in the current study, although the algorithm will also generalize to other clades. To identify such conserved proteins, we considered orthologous protein groups (orthogroups) on the hypothesis that orthologous proteins frequently retain similar domain compositions, at least within a limited taxonomic range, such as vertebrates (11, 12). This trend enables us to investigate whether rare domain compositions are nonetheless conserved across vertebrates within specific orthogroups. We thus developed our search method based on two criteria: {1} domains that rarely pair with transmembrane domains across vertebrate proteins, and {2} orthogroups in which the domain combination of interest occurs more frequently.

By focusing on rare but evolutionarily conserved combinations of transmembrane domains with transcription-related domains, we rediscovered well-known MBTFs, including SREBP, NOTCH, and ATF6, as well as other MBTRs that have been primarily studied for different biological functions. More importantly, we identified lesser-characterized transmembrane proteins with potential roles in gene regulation. One example is PREB (prolactin regulatory element-binding protein), which we experimentally validated as a potential MBTF. Our results suggested that PREB should undergo proteolytic cleavage, and in the absence of its transmembrane domain, PREB exhibited nuclear translocation and DNA binding activity. Additionally, we identified a potential MBTF class with the same DNA-binding domain as MYRF, which extends beyond vertebrates. We observed that this DNA-binding domain frequently pairs with transmembrane domains across vertebrates, prompting us to expand our analysis to all eukaryotic proteins that possess this domain, identifying a related but distinct candidate MBTR family in fungi. Our findings suggest that MBTRs could be more conserved than previously predicted, indicating their functional significance. This approach can be used to identify novel regulatory mechanisms across diverse biological contexts and taxonomic groups and can be applied to other protein domain combinations.

## Materials and methods

### Collecting domain annotations of vertebrate proteins and predicting transmembrane domains

We downloaded bulk protein data from UniProt (13) and collected Pfam domain annotations (8). To select vertebrate species and their subspecies, we downloaded the list of all descendants of vertebrates (Taxon ID 7742) from UniProt Taxonomy and chose taxa whose ranks were annotated as species or subspecies. Then, we retained proteins from these vertebrate species and subspecies. As a result, we selected 19,650,569 proteins. We ran TMbed (14) to predict whether these proteins have transmembrane domains. We excluded 1,599 proteins whose lengths exceeded 8,300 amino acids. We assigned 4,653,349 transmembrane proteins and 14,995,621 non-transmembrane proteins. In total, 99.992% of the selected vertebrate proteins were assigned.

### Mapping orthologous protein groups (orthogroups)

To define orthologous protein families, we used orthogroups from the eggNOG 5.0 database (15), assigning orthogroups to our protein sequences using eggNOG-mapper v2.1.9 (16). We filtered the mapping results using two criteria: first, we required that each protein have the same taxonomic annotation in UniProt and eggNOG, or that the UniProt taxon be a subspecies of the eggNOG species. We then removed orthogroups containing proteins from a single species only.

In total, we assigned 3,798,707 proteins from 102 species and their 92 subspecies to 27,745 orthogroups.

### Calculation of conservation-to-frequency ratios as a means to prioritize MBTRs

We find domains that rarely coexist with transmembrane domains by measuring the following frequency, referred to as 𝑓_𝑎𝑙𝑙_:

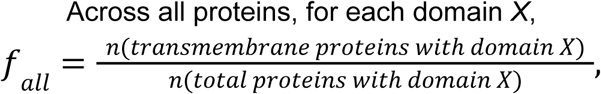

which was expected to be low, given our goal of identifying domains rarely found in a transmembrane context.

To identify domain combinations conserved across vertebrate species, we calculated the same ratio but for each orthogroup, referred to as 𝑓_𝑂𝐺_:

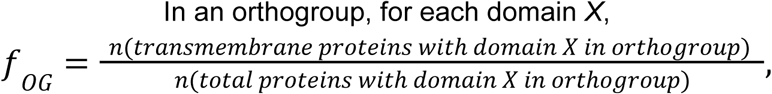

which was expected to be high for the case of well-conserved domain combinations.

Then, we divided 𝑓_𝑂𝐺_ by 𝑓_𝑎𝑙𝑙_, resulting in a list of conservation-to-frequency ratios (𝑓_𝑂𝐺_/𝑓_𝑎𝑙𝑙_), one for each [domain, orthogroup] pair. Domains that do not combine with transmembrane domains were excluded, and [domain, orthogroup] pairs with fewer than 10 transmembrane proteins were also filtered out. From the resulting distribution of conservation-to-frequency values, 1,239 [domain, orthogroup] pairs exhibited values exceeding 1.995 (10^0.3^). We selected 855 [domain, orthogroup] pairs that included at least one human or mouse protein for further analysis.

### Building phylogenetic trees with domain composition visualization

For SREBP1, NOTCH1, ATF6B, PREB, CERS2, and NAT8F3, we aligned their orthologous proteins (here, defined by assignment to the same pre-computed eggNOG orthogroup) using MAFFT v7.505 (17), and inferred approximately-maximum-likelihood phylogenetic trees using FastTree v2.1.11 (18). We selected the 10 most representative orthologs that have both transmembrane domains and domains of interest, using PARNAS v0.1.6 (19), prioritizing reviewed proteins (Swiss-Prot in UniProt). Pfam domains with their positions (location within a protein) were annotated using InterProScan v.5.75 (20). For visualizing phylogenetic trees and domains, we used the iTOL online tool (21). For proteins with “NDT80/PhoG-like DNA-binding family” domain and transmembrane domains, we downloaded sequences of all proteins with that domain annotation and selected proteins with transmembrane domains by using TMbed. We filtered out proteins with either the “MYRF ICA domain” or the “MYRF C-terminal domain 2” annotation from InterProScan. Subsequently, we constructed the phylogenetic tree and visualized it using iTOL as described above.

### Cloning of PREB and PREBΔTM

In order to create the modified construct pSBFB, the luciferase open reading frame (ORF) from the pSBtet-GP vector (Addgene, 60495) was replaced with a FLAG tag, a biotinylation site, and multiple cloning sites containing Notl and NheI restriction sites as previously described (22). Mouse *Preb* cDNA ORF clones were obtained from Sino Biological (MG5A1071-CY) and amplified using primers designed to produce constructs encoding PREB with or without the transmembrane domain (PREBΔTM) (**Supplementary Table S1**), with the incorporation of Notl and NheI restriction sites (NEB, R3189 and R3131). The amplified products were cloned into Notl/NheI-digested pSBFB vector using T4 DNA ligase (NEB, M0202). The cloned constructs were verified by Sanger sequencing and transfected into J1 mouse embryonic stem (ES) cells stably expressing BirA using Lipofectamine Stem Transfection Reagent (Thermo Fisher Scientific, STEM00008) along with pCMV(CAT)T7-SB100 (Addgene, 34879) for genomic integration. Transfected cells were selected with puromycin (1 µg/mL, Gibco A1113803) and G418 (250 µg/mL, Gibco 10131027) starting 24 hours after transfection to generate stable cell lines. To induce expression of PREB and PREBΔTM, cells were treated with 0.5 µg/mL of Doxycycline (Dox; Fisher BioReagents, BP26535) for 24 hours, and protein expression of both constructs was confirmed by western blotting.

### Cell culture

J1 mouse ES cells were cultured on plates coated with 0.1% gelatin in standard ES cell culture medium. The medium consisted of Dulbecco’s Modified Eagle’s Medium (DMEM; Gibco, 11965118), supplemented with 18% fetal bovine serum (FBS; MilliporeSigma, 12306C), EmbryoMax nucleosides (MilliporeSigma, ES-008-D), MEM non-essential amino acids (Gibco, 11-140-050), 0.1 mM β-mercaptoethanol (MilliporeSigma, M3148-25ML), 50 U/mL penicillin-streptomycin-glutamine (PSG, Gibco, 10378016), and 1000 U/mL recombinant mouse leukemia inhibitory factor (LIF; Gemini Bio-Products, 400-495). Cells were maintained at 37°C in a humidified incubator with 5% CO₂ and passaged every 2–3 days using 0.25% trypsin-EDTA (Thermo Fisher Scientific, 25200056) for single-cell dissociation.

### Western blot

PREB- and PREBΔTM-expressing mouse ES cells, treated with Dox for 24 hours, were lysed in Laemmli Sample Buffer (Bio-Rad, 1610747) supplemented with 5% β-mercaptoethanol (MilliporeSigma, M3148-25ML), then heated at 95°C for 10 minutes to denature proteins. Lysates were resolved by 10% SDS-PAGE, followed by transfer onto polyvinylidene fluoride (PVDF) membranes (MilliporeSigma, IPVH00010). Membranes were blocked with 5% skim milk in TBS-T (Tris-buffered saline containing 0.1% Tween-20) for 1 hour to prevent non-specific binding. Subsequently, the membranes were incubated overnight at 4°C with Streptavidin-HRP (Cytiva, RPN1231-2ML, 1:2000) to detect biotinylated PREB. β-actin antibody (Abgent, AM1829B, 1:20,000) was used as a loading control. After three 10-minute washes in TBS-T, membranes for β-actin detection were incubated with an anti-mouse HRP-conjugated secondary antibody (Cell Signaling Technology, 7076, 1:10,000) for 1 hour at room temperature. Following another set of three 10-minute washes with TBS-T, proteins were detected using enhanced chemiluminescence (ECL) reagent (Cytiva, 45-002-401).

### Immunofluorescence assays

PREB- and PREBΔTM-expressing mouse ES cells, with or without Dox treatment for 24 hours, were fixed in 4% paraformaldehyde for 20 minutes at room temperature, followed by three washes with PBS containing calcium and magnesium (PBS-CM; Corning, 21-040-CV) supplemented with 0.1% Bovine Serum Albumin (BSA; MilliporeSigma, A7906-500G). Following fixation, cells were permeabilized and blocked for 45 minutes at room temperature in PBS-CM/0.1% BSA containing 0.3% Triton X-100 (MilliporeSigma, T8787-50ML) and 10% horse serum (Gibco, 16050122). Cells were then incubated overnight at 4°C with Streptavidin conjugated to Alexa Fluor 555 (Thermo Fisher Scientific, S21381) in PBS-CM containing 0.1% BSA and 10% horse serum. The next day, cells were washed three times with PBS-CM/0.1% BSA, and nuclei were stained with 4′,6-diamidino-2-phenylindole (DAPI) for 10 minutes. After final washes, samples were imaged using a Nikon W1 Spinning Disk Confocal Microscope. Identical imaging settings were applied to all samples within each experiment to ensure consistency.

### bioChIP-sequencing experiment

bioChIP-seq was performed on PREB- and PREBΔTM-expressing mouse ES cells treated with 0.5 µg/mL Dox for 24 hours as previously described (23). Cells were cross-linked with 1% formaldehyde (Fisher Bioreagents, BP531-500) for 7 minutes at room temperature, and the reaction was quenched by adding glycine (Fisher Scientific, BP381-1) to a final concentration of 0.125 M. The chromatin was sheared to an average fragment size of ∼200 bp using a Bioruptor sonicator (Diagenode). Cross-linked chromatin was pre-cleared with Protein A agarose beads (Thermofisher, 10008D) and subsequently subjected to precipitation using Dynabeads MyOne Streptavidin T1 (Invitrogen, 65602). Precipitated complexes were washed twice with 2% SDS, once with buffer A (0.1% Deoxycholate, 1% Triton X-100, 1mM EDTA, 50mM HEPES pH 7.5, 500mM NaCl), once with buffer B (250mM LiCl, 0.5% NP-40, 0.5% Deoxycholate, 1mM EDTA, 10mM Tris-Cl pH 8.1), and twice with TE buffer. The crosslinked protein-DNA complexes in the elution buffer (1% SDS, 10mM EDTA, 50mM Tris-Cl pH 8) were incubated at 65°C overnight for reverse crosslinking. RNase A (Thermo Scientific, EN0531) was then treated for 30 minutes, followed by treatment of proteinase K (NEB, P8107S) for 2 hours. DNA was extracted with phenol:chloroform:isoamyl alcohol (Invitrogen, 15593031), then precipitated and re-suspended in the TE buffer. Library preparation was performed using the NEBNext Ultra DNA Library Prep Kit (NEB, 7103L), and sequencing was conducted on an Illumina NovaSeq 6000 platform to generate 50-bp paired-end reads. Comparison of the samples with the BirA-only control allowed identification of target-specific binding sites, and IgG ChIP served as an additional control.

### bioChIP-sequencing analysis

The ChIP-seq raw DNA sequencing data were adapter-trimmed using TrimGalore v0.6.10 (https://github.com/FelixKrueger/TrimGalore), aligned by Bowtie v2.5.1 (24), then processed with Samtools v1.20 (25) and Picard MarkDuplicates v3.0.0 (http://broadinstitute.github.io/picard). Peak calling was performed using Macs3 v3.0.1 (26), and the heatmap was generated using DeepTools v3.5.5 (27).

### Site-1 protease and Site-2 protease inhibitor treatment

PREB- and PREBΔTM-expressing mouse ES cells were seeded one day prior to treatment. The following day, cells were treated with 0.5 µg/mL Dox to induce expression, together with the S1P inhibitor PF-429242 (Selleck Chemicals, S6418) or the S2P inhibitor Nelfinavir Mesylate (Selleck Chemicals, S4282) at 10 µM. Cells were maintained under these conditions for 24 hours and 48 hours to assess the effects on proteolytic cleavage. DMSO was used as the control.

## Results

### A search for conserved domains that are found rarely in transmembrane proteins suggests new membrane-bound transcriptional regulators

In this study, we aimed to discover new MBTRs by developing a search method for protein domains that rarely pair with transmembrane domains, but, when found, are well-conserved across species, focusing here on vertebrate MBTRs. To develop the search method, we first collected information on vertebrate proteins, including their domain and orthogroup annotations, as well as the presence of transmembrane domains. We downloaded amino acid sequences and Pfam domain annotations (8) of 19.65 million (M) vertebrate proteins from UniProt (13). Transmembrane domains were predicted based on sequences using TMbed (14), resulting in 4.65M transmembrane proteins and 15M non-transmembrane proteins. To investigate evolutionary conservation, we transferred the eggNOG orthogroup annotations to our proteins. As a result, 27,745 orthogroups encompassing 3.8M proteins from 102 vertebrate species were assigned and used in the subsequent sections of this study (**Fig. 1**).

**Figure 1.**
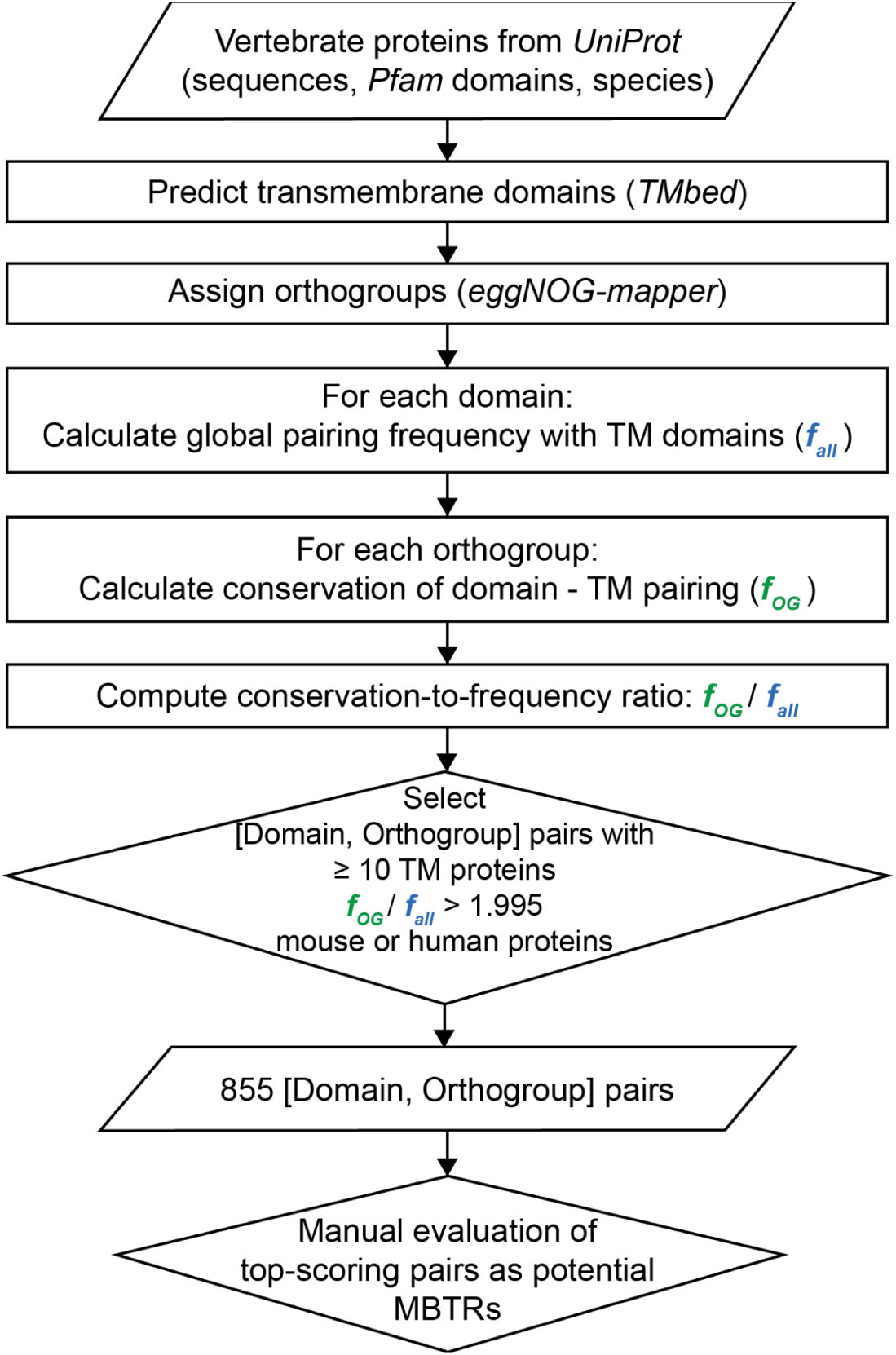
Search method to identify conserved protein domains that are rarely found in a transmembrane context as a means to discover potential new membrane-bound regulatory proteins.

To find domains that rarely pair with transmembrane domains, we calculated the frequency of transmembrane proteins for each domain across all proteins (𝑓_𝑎𝑙𝑙_; **Fig. 1**). If this value is low for a domain, it indicates that the domain and transmembrane domain rarely coexist in vertebrate proteins. Then, to check whether that domain combination is nonetheless conserved within specific orthogroups, we calculated the frequency of transmembrane proteins for each domain within each orthogroup (𝑓_𝑂𝐺_; **Fig. 1**). Since orthologous proteins can be highly conserved and can have highly similar domain combinations, the 𝑓_𝑂𝐺_ value can be correspondingly high in some orthogroups even if the overall 𝑓_𝑎𝑙𝑙_ is low. We then considered both the global frequency and the conservation of domain combinations by calculating the ratio of these frequencies (𝑓_𝑂𝐺_/𝑓_𝑎𝑙𝑙_; **Fig. 1**). We defined this ratio as the conservation-to-frequency value.

From this analysis, we made a list of all [domain, orthogroup] pairs with their corresponding conservation-to-frequency values. To remove spuriously large values coming from small orthogroups, we filtered out [domain, orthogroup] pairs with fewer than 10 transmembrane proteins. Then, based on the distribution of conservation-to-frequency values, we selected a heuristic cutoff (𝑓_𝑂𝐺_/𝑓_𝑎𝑙𝑙_= 1.995) for discovery (**Fig. 2**). Since there is no curated set of MBTRs, formal benchmarking (precision, recall, and FDR) against reference sets was not available. We excluded [domain, orthogroup] pairs that contained no human or mouse proteins, yielding 855 [domain, orthogroup] pairs (**Supplementary Table S2**), and examined the top-scoring families manually.

**Figure 2.**
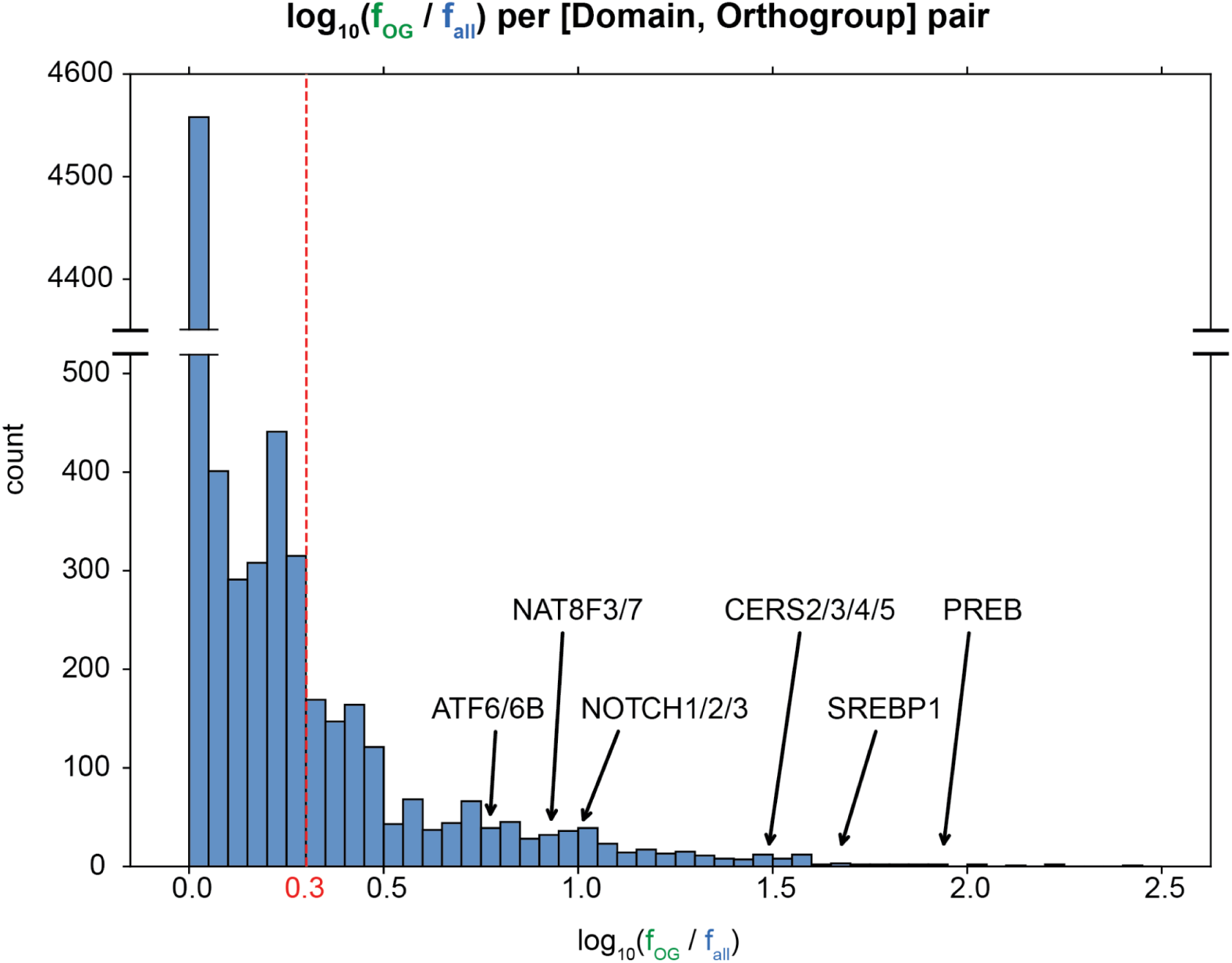
Known membrane-bound transcriptional regulators exhibit high conservation-to-frequency values (*f_OG_/f_all_*), accompanied by new candidate regulators.

Our method aims to find proteins satisfying the following two conditions: 1) having a domain that rarely pairs with transmembrane domains, but for which 2) the combination of domain of interest and transmembrane domain is nonetheless well-conserved in some orthogroups. The ratios corresponding to these two conditions were calculated and represented by 𝑓_𝑎𝑙𝑙_and 𝑓_𝑂𝐺_values, respectively. Then, we calculated the conservation-to-frequency values (𝑓_𝑂𝐺_/𝑓_𝑎𝑙𝑙_) for [domain, orthogroup] pairs. After filtering out [domain, orthogroup] pairs with annotated conditions, we investigated proteins in the final list.

The distribution of the logarithm base 10 of the conservation-to-frequency values for [domain, orthogroup] pairs with at least 10 transmembrane proteins. Based on this distribution, we selected a heuristic cutoff of 0.3. Three known MBTRs and three new candidate MBTRs are labeled.

### Well-known MBTFs were detected by our method

We first verified our method by assessing whether previously known MBTFs were caught. Our results included several well-known MBTFs: SREBP1 (Sterol regulatory element-binding protein 1), SREBP2, NOTCH1 (Neurogenic locus notch homolog protein 1), NOTCH2, NOTCH3, NOTCH4, ATF6 (Cyclic AMP-dependent transcription factor ATF-6 alpha), ATF6B, CREB3L1 (Cyclic AMP-responsive element-binding protein 3-like protein 1), CREB3L3, and CREB3L4 (**Table 1**). **Figure 3a** shows predicted structures of SREBP1, NOTCH1, and ATF6B from AlphaFold (28). The phylogenetic trees constructed from orthologs of these proteins illustrate the conservation of specific domain pairs across several vertebrate species (**Fig. 3b**). Both the conservation of domains at the orthogroup level and the AlphaFold structural predictions of alpha-helical transmembrane domains are in concordance with these proteins being correctly classified by the 𝑓_𝑂𝐺_/𝑓_𝑎𝑙𝑙_scores, suggesting this general workflow will also be valid for investigating subsequent new predictions.

**Figure 3.**
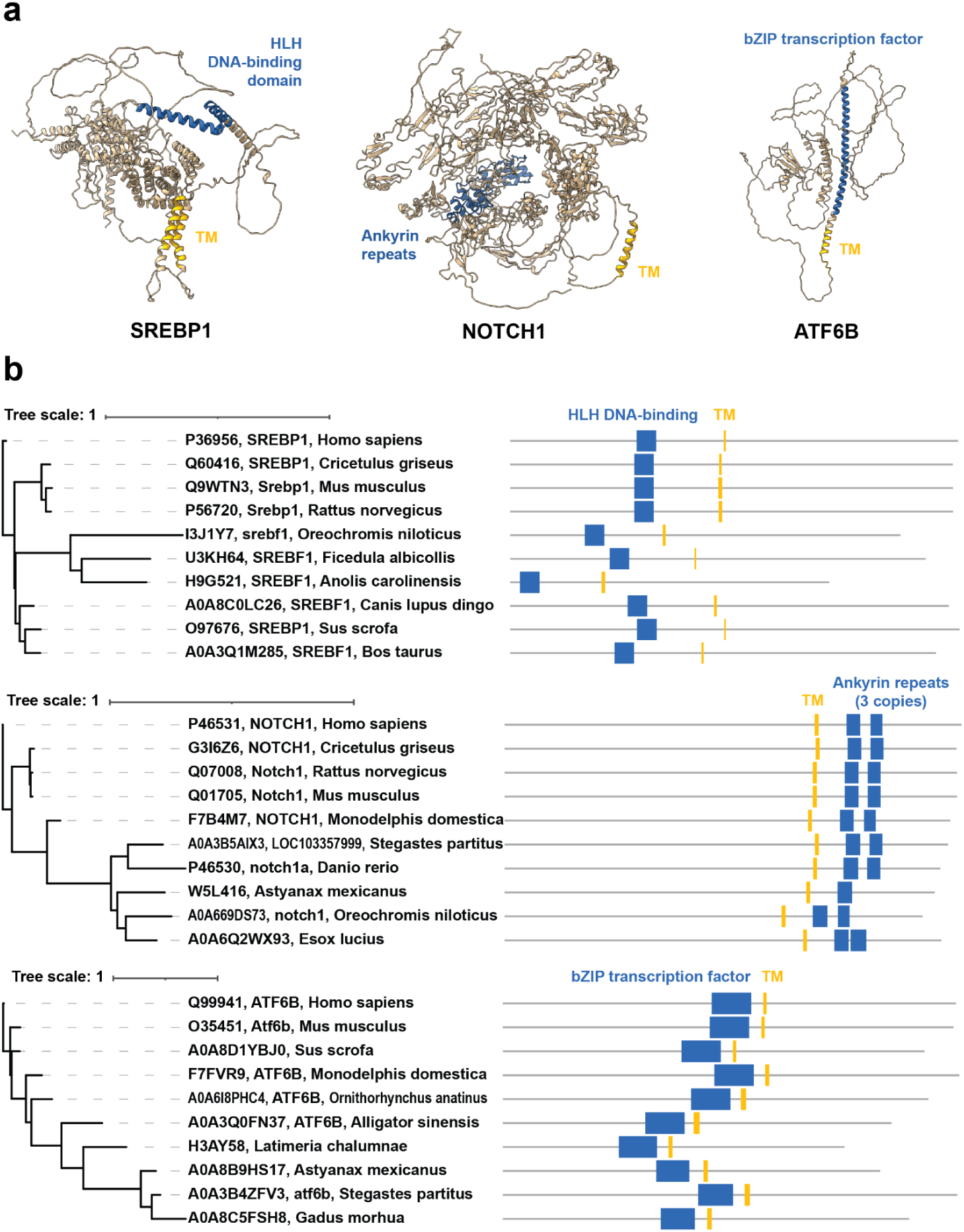
Predicted structures and phylogenetic trees of orthologs illustrate the conserved domain compositions for three well-known MBTFs with high conservation-to-frequency values. (**a**) AlphaFold predictions of three well-known MBTFs based on mouse protein sequences. Transmembrane domains are marked in yellow. The domains that comprise the combination with transmembrane domains are colored blue. (**b**) Phylogenetic trees showing 10 orthologs of each MBTF with their Pfam domain annotations (top: SREBP1, middle: NOTCH1, bottom: ATF6B). The same color scheme as panel (**a**) was applied.

**Table 1.** Well-known MBTFs detected by our method.

| Representative proteins | Pfam ID | Domain | $f_{all}$ | eggNOG ID | Orthogroup | $f_{OG}$ | $f_{OG}/f_{all}$ |
| --- | --- | --- | --- | --- | --- | --- | --- |
| ATF6 | PF00170 | bZIP transcription factor | 0.16 | ENOG50497SM | transcription factor | 0.96 | 5.93 |
| ATF6B | PF00170 | bZIP transcription factor | 0.16 | ENOG5048W9C | activating transcription factor 6 beta | 0.96 | 5.90 |
| CREB3L1 | PF00170 | bZIP transcription factor | 0.16 | ENOG50490X8 | cAMP responsive element binding protein 3-like 1 | 0.76 | 4.69 |
| CREB3L3 | PF00170 | bZIP transcription factor | 0.16 | ENOG50496Y8 | cAMP response element binding | 0.73 | 4.49 |
| CREB3L4 | PF00170 | bZIP transcription factor | 0.16 | ENOG5049MMU | cAMP response element binding | 1.00 | 6.18 |
|  | PF00170 | bZIP transcription factor | 0.16 | ENOG5049074 | cAMP response element binding | 0.73 | 4.53 |
| NOTCH1 | PF12796 | Ankyrin repeats (3 copies) | 0.10 | ENOG5048WDE | neurogenic locus notch homolog protein | 0.97 | 10.14 |
|  | PF07645 | Calcium-binding EGF domain | 0.35 | ENOG5048WDE | neurogenic locus notch homolog protein | 0.91 | 2.63 |
| NOTCH2 | PF13857 | Ankyrin repeats (many copies) | 0.09 | ENOG5048V3W | neurogenic locus notch homolog protein | 0.96 | 10.16 |
|  | PF12796 | Ankyrin repeats (3 copies) | 0.10 | ENOG50 48V3W | neurogenic locus notch homolog protein | 0.92 | 9.69 |
|  | PF07645 | Calcium-binding EGF domain | 0.35 | ENOG50 48V3W | neurogenic locus notch homolog protein | 0.80 | 2.28 |
| NOTCH3 | PF13857 | Ankyrin repeats (many copies) | 0.09 | ENOG50 49EBI | glomerular capillary formation | 0.96 | 10.22 |
|  | PF12796 | Ankyrin repeats (3 copies) | 0.10 | ENOG50 49EBI | glomerular capillary formation | 0.95 | 10.02 |
|  | PF07645 | Calcium-binding EGF domain | 0.35 | ENOG50 49EBI | glomerular capillary formation | 0.92 | 2.65 |
| NOTCH4 | PF00023 | Ankyrin repeat | 0.07 | ENOG50 497TI | Neurogenic locus notch homolog protein 4 | 0.99 | 14.32 |
|  | PF13637 | Ankyrin repeats (many copies) | 0.08 | ENOG50 497TI | Neurogenic locus notch homolog protein 4 | 0.96 | 12.12 |
|  | PF12796 | Ankyrin repeats (3 copies) | 0.10 | ENOG50 497TI | Neurogenic locus notch homolog protein 4 | 0.94 | 9.89 |
|  | PF07645 | Calcium-binding EGF domain | 0.35 | ENOG50 497TI | Neurogenic locus notch homolog protein 4 | 0.89 | 2.56 |
| SREBP1 | PF00010 | Helix-loop-helix DNA-binding domain | 0.01 | ENOG50 495AZ | sterol response element binding | 0.58 | 46.93 |
| SREBP2 | PF00010 | Helix-loop-helix DNA-binding domain | 0.01 | ENOG50 494W8 | sterol regulatory element binding transcription factor 2 | 0.16 | 12.74 |

SREBP1 had a conservation-to-frequency value of 46.93, ranking 10th overall, and SREBP2’s value, 12.74, also falls within the top 10%. Both proteins regulate lipid homeostasis and cholesterol synthesis (29), and their ‘helix-loop-helix DNA-binding domain’ was flagged by our approach. This domain exhibited a very low 𝑓_𝑎𝑙𝑙_value of 0.01, indicating that it rarely pairs with transmembrane domains, except across members of their corresponding ‘sterol response element binding’ and ‘sterol regulatory element binding transcription factor 2’ orthogroups. Interestingly, SREBP1 and SREBP2 had low 𝑓_𝑂𝐺_ values compared to other top-scoring protein families (0.58 and 0.16, respectively), but since their 𝑓_𝑎𝑙𝑙_ values were even lower, they scored well in our analysis. Thus, the combination of ‘Helix-loop-helix DNA-binding domain’ and transmembrane domains is somewhat less conserved, but this combination is even more scarce, and our approach is still able to assign a high score to this known MBTF family.

Different from SREBPs, all four NOTCH proteins were detected multiple times by different [domain, orthogroup] pairs with conservation-to-frequency values from 2.28 to 14.32. Our method flagged NOTCH proteins’ ‘ankyrin repeats’ and ‘calcium-binding EGF domain’ domains. The ankyrin repeat is one of the most common protein-protein interaction motifs and is found widely in proteins with diverse functions. The ‘calcium-binding EGF domain’ was named from the epidermal growth factor, where it was first described. This domain can be found in many membrane-bound and extracellular proteins (30), and our results showed that its 𝑓_𝑎𝑙𝑙_value is higher than that of ankyrin repeats. Orthogroups for NOTCH1, NOTCH2, and NOTCH4 were explicitly related to NOTCH homologs, while NOTCH3 belongs to a ‘glomerular capillary formation’ orthogroup, consistent with its known role in maintaining the structural integrity and functional stability of renal arteries (31). These orthogroups had high 𝑓_𝑂𝐺_ values, indicating that the combination of domains (of interest and transmembrane domain) was well conserved. Since NOTCH proteins are transmembrane receptors, this result was plausible.

The third protein group included ATF6 and ATF6B. Their conservation-to-frequency values were 5.93 and 5.90, respectively. These proteins are involved in the endoplasmic reticulum (ER) stress response, specifically the unfolded protein response. Under ER stress, the N-terminal cytoplasmic domain is cleaved and translocates to the nucleus (4, 32). The domain flagged by our method was ‘bZIP transcription factor,’ which has the basic region and the leucine zipper region. Its 𝑓_𝑎𝑙𝑙_value was 0.16, relatively low. Although the orthogroup of ATF6 is rather generically named ‘transcription factor’, almost all members of this orthogroup appear to correspond to ATF6 from various species. ATF6B’s orthogroup, ‘activating transcription factor 6 beta’, was also particular to ATF6B. These two orthogroups had high 𝑓_𝑂𝐺_ values, indicating that these proteins exhibit well-conserved, rare domain combinations.

The last protein group with the lowest conservation-to-frequency values among well-known MBTFs consisted of three CREB3L (Cyclic AMP-responsive element-binding protein 3-like) proteins: CREB3L1 (4.69), CREB3L3 (4.49), and CREB3L4 (6.18 and 4.53). CREB3L1 responds to ER stress or DNA damage, activating genes that inhibit cell cycle progression. CREB3L3 is proposed to act during ER stress and may function with ATF6. CREB3L4 may also contribute to the unfolded protein response (33–35). Our method flagged the ‘bZIP transcription factor’ domain, of which 𝑓_𝑎𝑙𝑙_value was 0.16. Orthogroups of these three proteins were ‘cAMP responsive element binding protein 3-like1’ and ‘cAMP response element binding’, and 𝑓_𝑂𝐺_ values were around 0.75, except for one small orthogroup (ENOG5049MMU). Those 𝑓_𝑂𝐺_values indicate that some CREB3L orthologous proteins lack the transmembrane domain.

### Our method rediscovered MBTRs that have been mainly studied for other biological functions

After validating that our search method can find well-known MBTFs, we asked if other less well-studied MBTFs, as well as the broader category of MBTRs, were scored highly by our search. In particular, we were interested in transmembrane proteins with evidence in the literature for regulating transcription, but for which this role was not necessarily well-characterized or broadly recognized. We identified nine MBTRs (**Table 2**; **Fig. 4**) that employ four distinct activation mechanisms, demonstrating the diversity of membrane-to-nucleus signaling. While their conservation-to-frequency values are generally modest compared to canonical MBTFs, their discovery underscores our method’s sensitivity in detecting regulatory proteins. (Since LRP1 is a large protein over 4,500 amino acids, its structure prediction is not included in **Figure 4**.)

**Figure 4.**
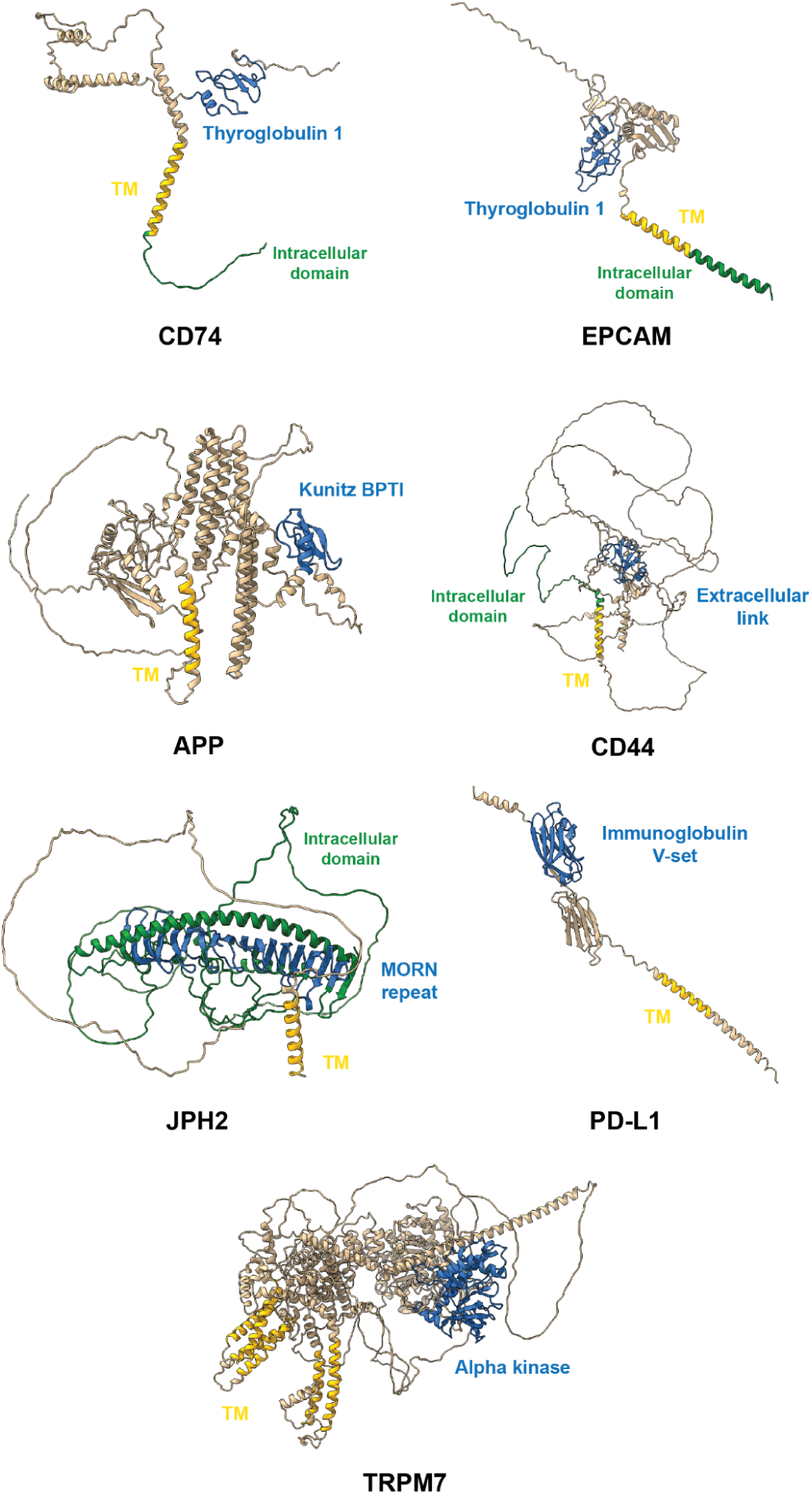
Predicted structures with domain annotations of MBTRs that have been mainly studied for their non-transcriptional-regulatory functions AlphaFold-predicted structures of MBTRs captured by our method that have primarily been studied for their non-transcriptional-regulatory biological functions. Transmembrane domains are marked in yellow, the domains that pair with transmembrane domains are marked blue, and cytoplasmic intracellular domains are colored in green if available. (The structure of LRP1 was excluded due to its massive size, exceeding 4,500 amino acids.)

**Table 2.** Characterized MBTRs rediscovered by our search that have been mainly studied for other biological functions.

| Category | Representative proteins | Pfam ID | Domain | $f_{all}$ | eggNOG ID | Orthogroup | $f_{OG}$ | $\frac{f_{OG}}{f_{all}}$ |
| --- | --- | --- | --- | --- | --- | --- | --- | --- |
| Regulated Intramembrane Proteolysis (RIP)-dependent | CD74 | PF00086 | Thyroglobulin type-1 repeat | 0.13 | ENOG50494AH | MHC class II protein binding, via antigen binding groove | 0.98 | 7.56 |
|  | EPCAM | PF00086 | Thyroglobulin type-1 repeat | 0.13 | ENOG50491FU | Epithelial cell adhesion molecule | 0.96 | 7.39 |
|  | LRP1 | PF12662 | Complement C1r-like EGF-like | 0.19 | ENOG5048WNX | alpha-2 macroglobulin receptor activity | 0.77 | 4.06 |
|  |  | PF07645 | Calcium-binding EGF domain | 0.35 | ENOG5048WNX | alpha-2 macroglobulin receptor activity | 0.80 | 2.30 |
|  | APP | PF00014 | Kunitz/Bovine pancreatic trypsin inhibitor domain | 0.31 | ENOG50 494FN | Amyloid beta (A4) | 0.93 | 3.03 |
|  | CD44 | PF00193 | Extracellular link domain | 0.38 | ENOG50 4993T | CD44 molecule (Indian blood group) | 0.91 | 2.40 |
| Stress-responsive proteolytic release | JPH2 | PF02493 | MORN repeat | 0.16 | ENOG50 490UH | phosphatidylinositol-5-phosphate binding | 0.62 | 3.73 |
| Non-proteolytic nuclear translocation | PD-L1 (CD274) | PF07686 | Immunoglobulin V-set domain | 0.32 | ENOG50 48WVQ | Programmed cell death 1 ligand | 0.86 | 2.66 |
| Enzymatically active domain release | TRPM6, TRPM7 | PF02816 | Alpha-kinase family | 0.39 | ENOG50 48X4Y | calcium-dependent cell-matrix adhesion | 0.97 | 2.48 |

The largest group (5 proteins: CD74, EPCAM, LRP1, APP, and CD44) is activated through regulated intramembrane proteolysis (RIP), a conserved mechanism in which sequential proteolytic cleavages release an intracellular domain (ICD) that translocates to the nucleus (36–43). CD74 and EPCAM exhibit the highest conservation-to-frequency values (7.56 and 7.39) among the nine rediscovered MBTRs. CD74 releases an intracellular domain (CD74-ICD) that modulates genes involved in apoptosis, immune response, and cell migration through interactions with RUNX and NF-κB (37). Similarly, the cleavage of EPCAM generates the EPCAM intracellular domain (EP-ICD), which drives genes involved in cell division, pluripotency, and EMT-associated processes (44). LRP1, APP, and CD44 showed lower values (4.06/2.30, 3.03, and 2.40) but share a similar architecture. LRP1-ICD co-activates PPARγ, a central regulator of lipid and glucose metabolism (45). APP’s ICD regulates multiple genes, including KAI1, Neprilysin, and p53 (41). CD44-ICD activates matrix metalloproteinases (46).

The second activation mechanism employs a stress-induced proteolytic cleavage, exemplified by JPH2, a membrane protein cleaved in cardiomyocytes under mechanical stress. This stress-induced cleavage generates an N-terminal fragment that translocates to the nucleus and activates cardioprotective genes, demonstrating how structural and transcriptional reprogramming are coordinated in the stressed heart (47). PD-L1 demonstrates an alternative activation mechanism. Although it is primarily observed in cancer cells, deacetylation of PD-L1 by HDAC2 triggers its nuclear translocation without cleavage, where it regulates immune-response genes (48). This case demonstrates that the membrane anchoring can be controlled epigenetically, not always through irreversible proteolytic cleavage. TRPM6 and TRPM7 (conservation-to-frequency value: 2.48) expand the MBTRs’ functional scope beyond TFs. They are ion channels but contain a serine/threonine kinase at their carboxyl termini. Upon cleavage, their C-terminal kinase domains translocate to the nucleus and phosphorylate histones, thereby directly modifying chromatin structure (49–51).

Together, these nine proteins illustrate the mechanistic diversity underlying membrane-bound transcriptional regulation, encompassing canonical RIP, stress-induced cleavage, non-proteolytic nuclear translocation, and cleavage-dependent release of histone kinase. Evolution has thus repeatedly used switchable membrane anchoring as a key regulatory component across a variety of biological processes. Importantly, these examples illustrate that our approach is agnostic as to the specific mechanism employed or the particular domains used, hence opening the possibility for discovering new classes of MBTRs.

### New candidate MBTRs include PREB, ceramide synthases, and N-acetyltransferases

Having confirmed that our search method could identify both already-known MBTFs and MBTRs more generally, we next searched for novel candidate MBTRs amongst the high-scoring protein families (**Table 3**; **Fig. 5**).

**Figure 5.**
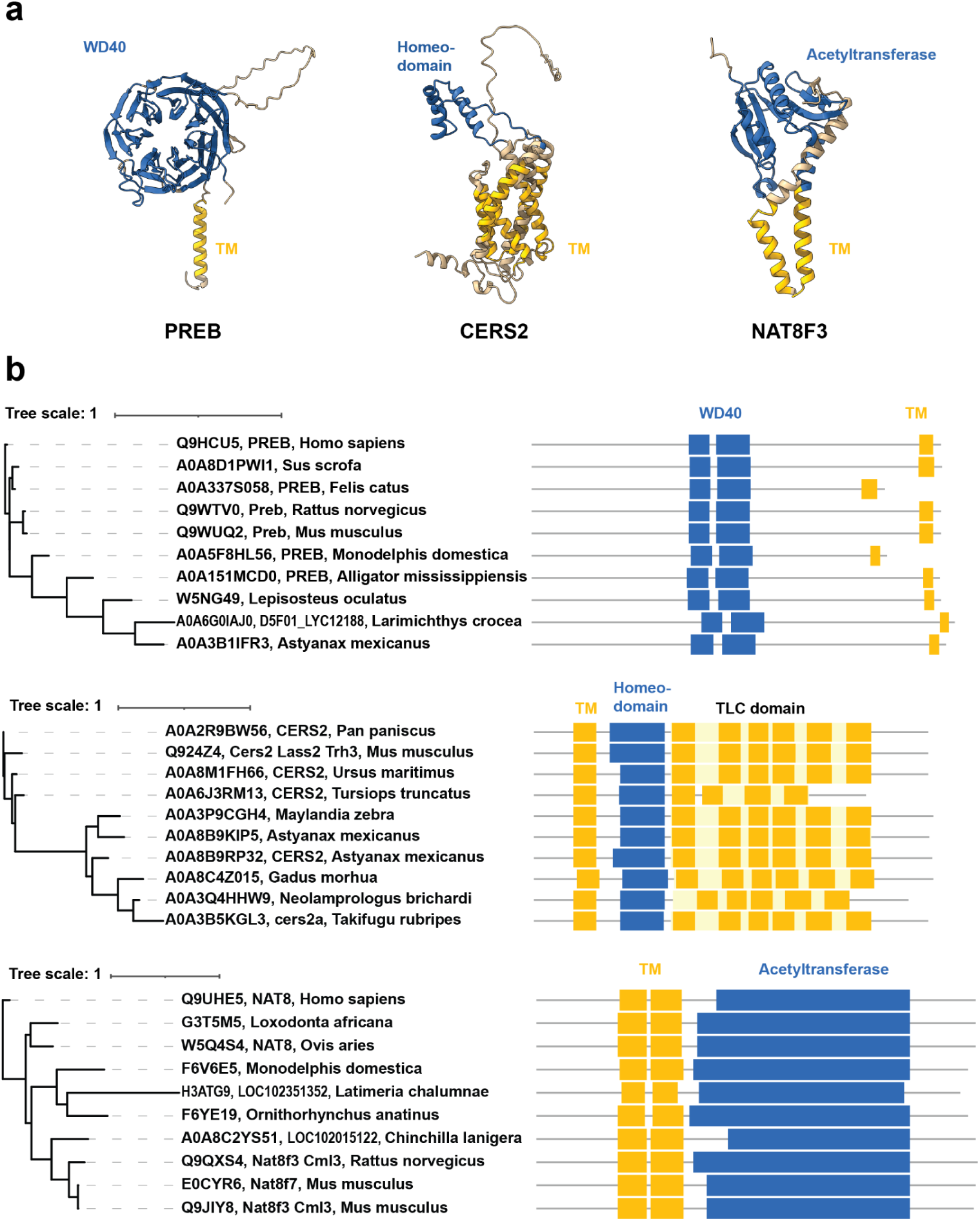
Based on their conserved domain compositions, PREB, CERS2, and NAT8F3 are new candidate MBTRs (**a**) AlphaFold-predicted structures for three potential MBTRs, here showing the mouse proteins. Transmembrane domains are marked in yellow. The domains that comprise the combination with the transmembrane domain are colored blue. (**b**) Phylogenetic trees showing 10 orthologs of each candidate protein with their Pfam domain annotations (top: PREB, middle: CERS2, bottom: NAT8F3 and NAT8F7). The TLC domain includes internal transmembrane domains. The same color scheme as panel (**a**) was applied.

**Table 3.** New candidate MBTRs.

| Representative proteins | Pfam ID | Domain | $f_{all}$ | eggNOG ID | Orthogroup | $f_{OG}$ | $f_{OG}/f_{all}$ |
| --- | --- | --- | --- | --- | --- | --- | --- |
| PREB | PF00400 | WD domain, G-beta repeat | 0.01 | ENOG504999H | prolactin regulatory | 0.84 | 86.57 |
| CERS2 | PF00046 | Homeodomain | 0.03 | ENOG5048YJT | regulation of Schwann cell proliferation involved in axon regeneration | 1.00 | 30.76 |
| CERS3 | PF00046 | Homeodomain | 0.03 | ENOG50497XU | sphingosine N-acyltransferase activity | 1.00 | 30.76 |
| CERS4 | PF00046 | Homeodomain | 0.03 | ENOG5048X03 | sphingosine N-acyltransferase activity | 1.00 | 30.76 |
| CERS5 | PF00046 | Homeodomain | 0.03 | ENOG5048VWA | sphingosine N-acyltransferase activity | 1.00 | 30.76 |
| NAT8F3, NAT8F7 | PF00583 | Acetyltransferase (GNAT) family | 0.12 | ENOG5049MJF | N-acetyltransferase | 0.98 | 8.50 |

PREB (prolactin regulatory element-binding protein) was ranked one of the highest-scoring protein families of our entire analysis. It has a ‘WD domain, G-beta repeat’ domain annotation with the 𝑓_𝑎𝑙𝑙_ value 0.01, one of the lowest values in the results. WD repeat domains are known to interact with diverse binding partners, such as proteins, DNA, and RNA (52). PREB’s WD repeat domain forms a seven-bladed beta-propeller structure, with a short alpha-helix transmembrane domain on the C-terminal side (**Fig. 5a**). PREB’s orthogroup annotation was ‘prolactin regulatory’ with an 𝑓_𝑂𝐺_value of 0.84, mostly consisting of PREB proteins from various species. The phylogenetic tree of 10 PREB orthologs shows conservation of the WD and transmembrane domains across vertebrates (**Fig. 5b**).

The next most promising candidates in our results were ceramide synthases (CERSs). CERS2, CERS3, CERS4, CERS5, and CERS6, which have multiple transmembrane domains. Ceramide synthases catalyze the transfer of the acyl chain from acyl-CoA to a sphingoid base, and six isoforms of mammalian CERS, except CERS1, contain a homeodomain (53). Our results recovered their homeodomain annotation (PF00046; 𝑓_𝑎𝑙𝑙_: 0.03; **Fig. 5a**). Their orthogroups (‘sphingosine N-acyltransferase activity’, ‘regulation of Schwann cell proliferation involved in axon regeneration’, 𝑓_𝑂𝐺_ values were 1.00 for both) were highly CERS-specific. It was reported that Schlank, a Drosophila CERS, binds to DNA via its homeodomain, and mouse CERS2 can also regulate transcription (54). However, human CERS2 lacks residues critical for DNA binding, making it unclear whether the homeodomain’s binding function has been conserved. Still, the phylogenetic tree of 10 CERS2 orthologs shows the conservation of both the homeodomain and transmembrane domains across several vertebrate species (**Fig. 5b**).

Another interesting case is NAT8F3 (N-acetyltransferase family 8 member 3) and NAT8F7, mouse proteins reported to be histone acetyltransferases that prefer histone H4 and exhibit perinuclear localization. It was suggested that these proteins are conserved across chordates but originated in cnidaria (55). Their domain annotation was ‘acetyltransferase (GNAT) family’ (PF00583; 𝑓_𝑎𝑙𝑙_: 0.12; **Fig. 5a**), and the orthogroup was ‘N-acetyltransferase’ (𝑓_𝑂𝐺_: 0.98). The phylogenetic tree of 10 orthologs shows both acetyltransferase domains and transmembrane domains (**Fig. 5b**).

While the extent to which these histone acetyltransferase activities are conserved across vertebrates is currently unknown, our analysis suggests that this family could be a new MBTR, along with PREB and the ceramide synthases. In order to provide more direct experimental support for this activity, we next focused our effort on evaluating the top-scoring PREB family.

### PREB without its transmembrane domain suggests a potential processing event, nuclear translocation, and DNA binding

PREB was historically identified as a DNA-binding protein that regulates prolactin, a hormone responsible for lactation (56). It was subsequently reported that PREB also binds to the promoters of insulin and adiponectin, and regulates gluconeogenic genes (57–59). Therefore, PREB’s function appeared from these data to be predominantly related to lactation, glucose, and lipid metabolism. However, there are two apparently largely disjoint sets of literature observations on PREB, which is also alternatively known as Sec12, a guanine nucleotide exchange factor that plays a crucial role in the formation of coat protein complex II (COPII) vesicles (60) and, moreover, is far more widespread in expression than its historic name would suggest. COPII vesicles transport proteins from the ER to the Golgi apparatus. For this function, PREB should be anchored on the membrane. In contrast, as a TF, PREB should enter the nucleus. These two bodies of literature suggest that the PREB protein has a dual functionality. Considering PREB as a candidate MBTR would thus imply that the membrane-bound form is responsible for regulating vesicle formation, while a proteolytically-released cytoplasmic domain might translocate to the nucleus to regulate transcription.

To investigate whether PREB lacking its transmembrane domain (PREBΔTM) translocates to the nucleus and binds DNA, we ectopically expressed cDNAs for full-length PREB and PREBΔTM in mouse embryonic stem (ES) cells using the pSBFB doxycycline (Dox)-inducible overexpression system (23, 61, 62) (**Fig. 6a,b**). Dox induction did not elicit detectable morphological changes for both cases (**Supplementary Fig. S1**). However, Western blot analysis revealed multiple protein bands following induction of full-length PREB, whereas PREBΔTM appeared as a single band (**Fig. 6c**), suggesting potential proteolytic processing of the full-length protein. Notably, as the expression clone was derived from a cDNA ORF, we can rule out alternative splicing in this context. Because the pSBFB vector introduces an N-terminal tag detectable by streptavidin-HRP, the presence of the tag in both bands indicates that processing occurs at the C-terminus (**Fig. 6c**).

**Figure 6.**
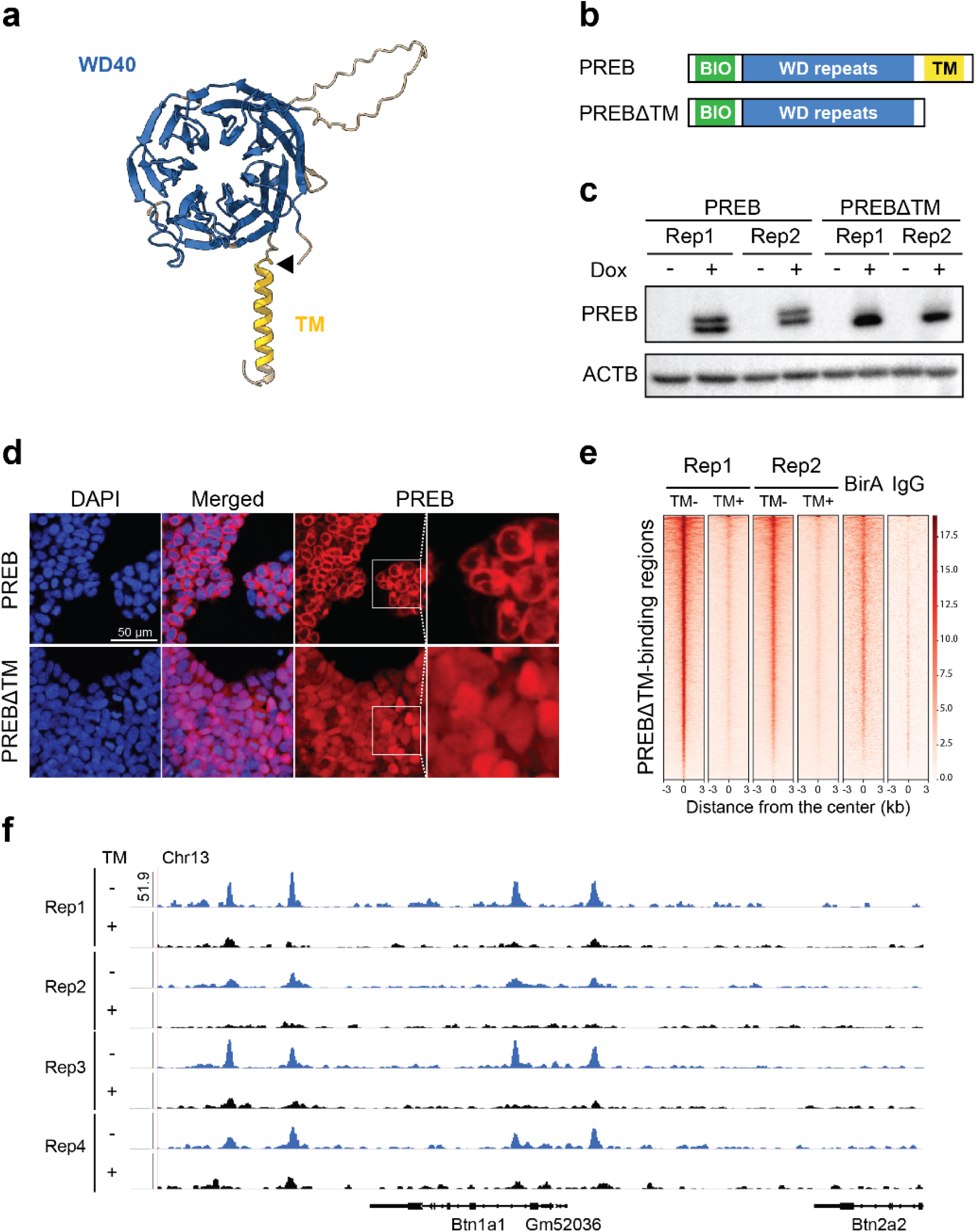
PREB without its transmembrane domain undergoes nuclear translocation and binds to the butyrophilin locus. (**a**) Predicted structure of PREB generated by AlphaFold, highlighting the WD repeat domains, the transmembrane domain, and the site of an engineered C-terminal truncation. (**b**) Schematic of the two constructs created to assay the mouse full-length PREB protein and PREBΔTM, lacking the C-terminal transmembrane domain. The N-terminal biotinylation site is indicated. (**c**) Western blot results of two biological replicates of ES cells expressing PREB and PREBΔTM. Two bands for PREB were detected by Streptavidin-HRP, suggesting C-terminal processing of a full-length PREB when expressed in mouse ES cells. (**d**) Immunofluorescence confocal microscopy images of PREB and PREBΔTM following Dox induction. PREB localizes predominantly to a perinuclear region, forming a ring-like shape. PREBΔTM displays diffused nuclear localization. Rightmost panels show zoomed images corresponding to the boxed regions. (**e**) bioChIP-seq profiles for PREB, PREBΔTM, BirA-only Strep-pulldown, and IgG ChIP-seq control samples. PREBΔTM samples show substantially stronger DNA-binding activity than full-length PREB. (**f**) Representative locus-specific binding profiles of PREB and PREBΔTM at the Btn1a1 (butyrophilin subfamily 1 member A1) gene. PREBΔTM shows markedly higher peak intensities, consistent with enhanced nuclear localization in the absence of the transmembrane domain.

Immunofluorescence imaging showed a perinuclear ring for full-length PREB consistent with ER proximity, while PREBΔTM displayed a predominant nuclear signal by DAPI overlap (**Fig. 6d**), supporting the hypothesis that the transmembrane domain prevents nuclear translocation. Given the nuclear localization of PREBΔTM, we hypothesized that it may exhibit stronger DNA-binding activity than PREB. To test this, we performed biotin-mediated chromatin immunoprecipitation followed by sequencing (bioChIP-seq) upon ectopic expression of PREB and PREBΔTM. To confirm the specificity of the binding events, we used a BirA-only control to check endogenous biotinylation and an IgG control for non-specific association. Aligning with the nuclear localization signal of PREBΔTM, the bioChIP-seq result showed that PREBΔTM exhibited a markedly greater increase in genome-wide DNA binding compared with full-length PREB (**Fig. 6e**). Based on previous literature (56), we expected PREBΔTM binding peaks near the prolactin gene, but this was not observed in ES cells under the conditions tested. We detected the binding peaks near other genes, most notably the butyrophilin gene (**Fig. 6f**).

Although these experiments do not reflect physiological conditions, the results indicate that PREBΔTM can relocate into the nucleus and associate with chromatin, supporting PREB as a candidate MBTF, pending validation of endogenous processing in relevant contexts. We note that immunohistochemistry of endogenous PREB in human appendix, colon, and rectum tissues from Human Protein Atlas (63) does show that PREB can natively localize to the nucleus of a subset of cells in these tissues (**Supplementary Fig. S2)**; unfortunately, no such data exist for the pituitary gland, where PREB was first isolated as a DNA binding protein, although experiments by Suzuki Fliss and co-authors confirmed nuclear localization in the rat pituitary tumor GH3 cell line (56).

Since the proteases responsible for PREB cleavage in ES cells and the precise cleavage site remain unknown, we also investigated whether PREB has a canonical site-1 protease (S1P) motif, a feature shared with multiple other known MBTFs (64), and found matched sequences. Since site-2 protease (S2P) acts in concert with S1P, we treated PREB-expressing ES cells with inhibitors of S1P and S2P (PF-429242 and Nelfinavir, respectively (65, 66). However, at least under the conditions tested, inhibition of these proteases did not detectably alter PREB cleavage (**Supplementary Fig. S3)**.

### Applying the method to a broader range of species, we found a potential MBTF family

Our method found known and novel potential MBTRs, but protein myelin regulatory factor (MYRF), a well-known MBTF, wasn’t rediscovered. We investigated MYRF’s domains and found that, in our method, MYRF scored poorly because its domains frequently pair with transmembrane domains in vertebrates. Notably, one of the domains is a known DNA-binding domain, ‘NDT80/PhoG-like DNA-binding family,’ related to the yeast meiosis-specific transcription factor NDT (67). We wondered whether this specific domain paired frequently with transmembrane domains outside of vertebrates, so we applied our method more broadly to eukaryotic proteins having the ‘NDT80/PhoG-like DNA-binding family’ annotation. As a result, we found three orthogroups with high 𝑓_𝑂𝐺_values. Two orthogroups contained MYRF and corresponded to MYRF orthologs, with some occurring outside of animals, as for the choanoflagellate *Monosiga* (e.g., the gene A9UP12_MONBE), which lacks myelin. This broad phylogenetic distribution of MYRF orthologs has been well-characterized previously, with MYRF family members pre-dating animals and controlling transcriptional programs unrelated to myelin in these other species, as previously reviewed (68).

However, in addition, we also identified a small protein orthogroup of unknown function (ENOG502D42H) that does not include MYRF, and which may represent a new fungal MBTF class comprising the NDT80 DNA-binding domain with a conserved transmembrane domain, but which lacks MYRF’s signature intramolecular chaperone autoprocessing (ICA) domain (**Fig. 7a**; see the first five proteins marked with a vertical bar in **Figure 7b**). We investigated whether there are more proteins outside of this orthogroup that have both the “NDT80/PhoG-like DNA-binding family” and transmembrane domains, without other typical MYRF-related protein domains (“Myelin regulatory factor ICA domain” and “Myelin gene regulatory factor C-terminal domain 2”) (**Fig. 7b**). The result includes two new fungal proteins with the same domain composition as the aforementioned small subfamily, which could be additional homologs. Other proteins have additional unrelated domains: Four proteins from the *Fusarium* genus share the same “Protein of unknown function (DUF3176)” domain. Seven proteins from various species (two fish species, one sponge, and four species belonging to Amoebozoa) contain a “Chaperone of endosialidase” domain, which is also found in MYRF, but they lack MYRF’s C-terminal domain. Based on the variation even in this one family, we strongly suspect that applying our method to a broader range of species than vertebrates should enable the identification of considerably more MBTRs.

**Figure 7.**
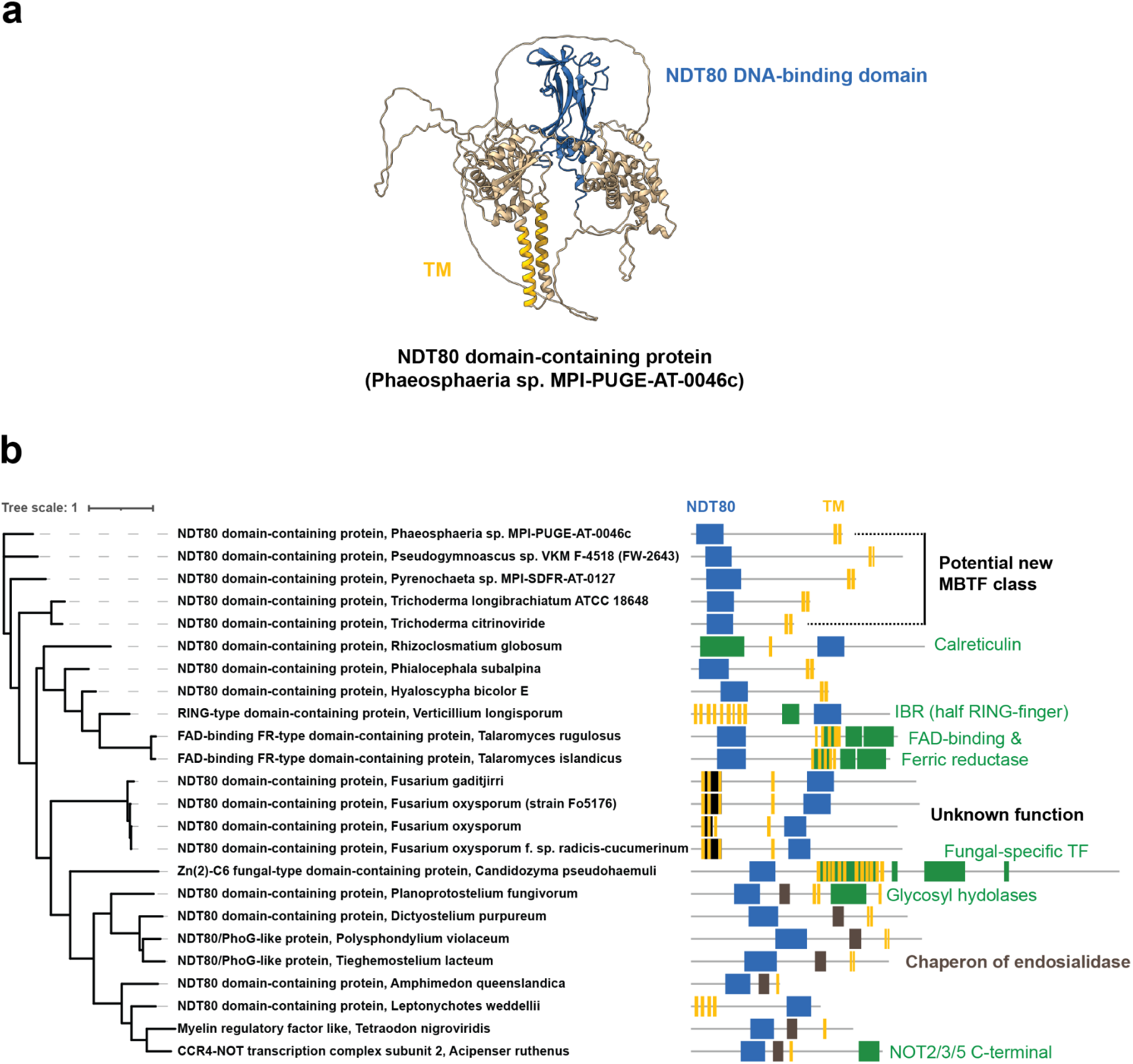
A predicted structure and domain compositions of potential MBTF orthologs containing the NDT80 DNA-binding domain (**a**) AlphaFold-predicted structure of an NDT80 domain-containing protein (UniProt entry: A0A9P9Q0Z2_9PLEO). Predicted transmembrane domains are marked in yellow, and the NDT80 DNA-binding domains are shown in blue. (**b**) Phylogenetic tree and domain compositions of eukaryotic proteins harboring both the “NDT80/PhoG-like DNA-binding family” domain and transmembrane domains, but lacking MYRF-related domains (“Myelin regulatory factor ICA domain” and “Myelin gene regulatory factor C-terminal domain 2”). The same color scheme as in panel (**a**) is used, with additional colors (green, brown, and black) for other protein domains. The vertical bar denotes a putative novel fungal MBTF family.

## Discussion

In this study, we developed a novel strategy to identify MBTRs based on protein domain combinations and conserved orthogroups. This approach successfully rediscovered known MBTFs and membrane-bound chromatin modifiers, while also revealing candidate MBTRs that had not been previously recognized. We further observed that several MBTFs were initially characterized for biological roles unrelated to transcriptional regulation, with their regulatory roles recognized only later. This trend underscores the utility of our method for identifying multifunctional proteins whose atypical modes of transcriptional regulation have been previously overlooked. Moreover, our findings suggest that MBTRs are conserved across vertebrate species and are even more broadly distributed across eukaryotes, as exemplified by NDT80 DNA-binding domain proteins. These results are consistent with a widespread regulatory paradigm in which extracellular cues are coupled to transcriptional responses within well-represented orthogroups. Quantitative assessment of this prevalence will require expanded orthology resources, systematic sensitivity analyses, and functional validation across diverse taxonomic groups.

Our identification of diverse MBTRs reveals that transcriptional regulation in membrane-bound contexts operates through multiple mechanisms that extend beyond the canonical regulated intramembrane proteolysis (RIP) observed in proteins such as SREBP and ATF6 (3). While RIP represents the predominant activation mode for MBTRs we identified, proteins such as PD-L1 demonstrate that membrane anchoring can also be modulated by post-translational modifications, including deacetylation, thereby allowing nuclear translocation without irreversible proteolytic cleavage. The presence of non-proteolytic mechanisms suggests that certain MBTRs may function as reversible transcriptional switches. Furthermore, membrane-bound histone modifiers, such as TRPM6/7, whose kinase domains modify chromatin structure via histone phosphorylation, expand the functional repertoire of MBTRs beyond classical TFs. Collectively, the mechanistic diversity highlights membrane anchoring as a versatile regulatory strategy, providing a spatial sequestration of transcriptional regulators until appropriate signals trigger their release.

Nevertheless, several considerations apply to our search strategy. First, our domain-based framework naturally depends on current domain annotations and transmembrane prediction tools. Second, TMbed, Pfam, and eggNOG were selected for their scalability, and the successful recovery of NOTCH, SREBP, and ATF6 supports their suitability for this screen. Third, when we narrow down [domain, orthogroup] pairs, we applied filters to reduce noise in conservation-to-frequency ratios; however, these criteria may preferentially enrich for larger, well-annotated protein families. Recovery of SREBP1/2, NOTCH 1-4, and ATF6/6B at high conservation-to-frequency ratio scores supports the utility of our strategy. Nonetheless, candidate rankings can shift with the threshold settings, minimum transmembrane protein filters, and species sampling.

We validated PREB as a candidate MBTF with dual functionality by demonstrating that deletion of its transmembrane domain alters its subcellular localization and enables DNA binding, thereby mimicking the outcome of a potential proteolytic activation mechanism. Future proteomic approaches to identify interaction partners and post-translational modifications will be essential for elucidating the mechanisms underlying its dual roles. This behavior supports a model in which membrane anchoring functions as a molecular switch that controls the spatial and temporal deployment of PREB activity. Such dual functionality might be more widespread than we have acknowledged. The evolution of such bifunctional proteins, which serve as both membrane-bound sensors and transcriptional regulators, represents a simple yet effective solution for coordinating multiple biological processes across cellular compartments. The molecular architecture of MBTRs may offer a practical framework for engineering synthetic regulatory switches that link membrane-associated signals to programmable gene expression responses.

While our *in vitro* experiments with PREB provide compelling evidence for its function as a potential MBTF that undergoes proteolytic cleavage, several limitations warrant consideration. Our studies were conducted in mouse ES cells, which may not reflect the physiological context in which PREB exerts its native functions. Previous studies have argued that PREB plays prominent roles in lactation, glucose metabolism, and lipid metabolism, implicating tissues such as the mammary glands and other metabolically active organs as more relevant settings for functional assessments. A visual survey of PREB localization as measured by immunohistochemistry in the Human Protein Atlas (63) reveals mixed cytoplasmic/membrane localization in many cell types assayed, but strong nuclear localization in a subset of cell types, as in the human appendix, colon, and rectum tissues in **Supplementary Fig. S2**. At this stage, we cannot rule out alternative splicing of the endogenous gene as a means of controlling differential localization, which will require detailed characterization of splice isoforms in cell types showing the differential localization. ChIP-seq experiments in these tissues will also be important for identifying context-specific target genes and refining PREB’s biological roles. Additionally, although an overexpression-based system has been shown to be effective, it may perturb endogenous regulatory mechanisms. Lastly, while the presence of multiple bands in the western blot suggests proteolytic processing, we did not directly map cleavage products, and alternative post-translational modifications or partial degradation cannot be excluded at this stage.

Finally, although our study focused on proteins containing transmembrane domains in combination with other domains, especially DNA-relevant domains, to identify transmembrane transcriptional regulators, the underlying strategy is readily extensible to other types of regulatory proteins characterized by rare domain combinations. For example, proteins that couple RNA-binding domains with enzymatic domains might be candidates to link metabolic signals and post-transcriptional regulation, whereas metabolic enzymes fused to cytoskeletal proteins could integrate metabolic feedback regulation to cell structural form or integrity. More broadly, our orthogroup-based filtering strategy can also be applied to any domain combination of interest to assess whether such domain combinations are preserved intact over evolution and thus likely to be functionally meaningful. In this way, the approach provides a generalizable way to identify multifunctional proteins that coordinate diverse cellular processes through unexpected combinations of protein domains.

## Acknowledgements

The authors acknowledge the Texas Advanced Computing Center at The University of Texas at Austin for providing high-performance computing resources that contributed to the research results reported in this paper. We also thank the Center for Biomedical Research Computing Facility (BRCF) PODs for computational support in data analysis. Some elements of the graphical abstract were created with BioRender.com (https://biorender.com/ftdvmzg).

## Author contributions

Muyoung Lee (Conceptualization, Data analysis, Writing—original draft, Writing—review and editing), Yeejin Jang (Performing experiments, Writing—original draft, Writing—review and editing), Qingqing Guo (Performing experiments), Jonghwan Kim (Supervision, Writing—original draft, Writing—review and editing), and Edward M. Marcotte (Conceptualization, Supervision, Writing—original draft, Writing—review and editing).

## Supplementary data

Supplementary data are appended at the end of the manuscript. The table of 855 [domain, orthogroup] pairs and proteins belonging to each case is available as Supplementary Table 2 and can be downloaded from the supporting Zenodo repository.

## Data availability

The ChIP-seq data have been submitted to the NCBI Gene Expression Omnibus (GEO; https://www.ncbi.nlm.nih.gov/geo/) under accession number GSE309766.

## Abbreviations

TF: transcription factor
MBTF: membrane-bound transcription factor
MBTR: membrane-bound transcriptional regulator

## Supplementary Figures

**Supplementary Figure S1.**
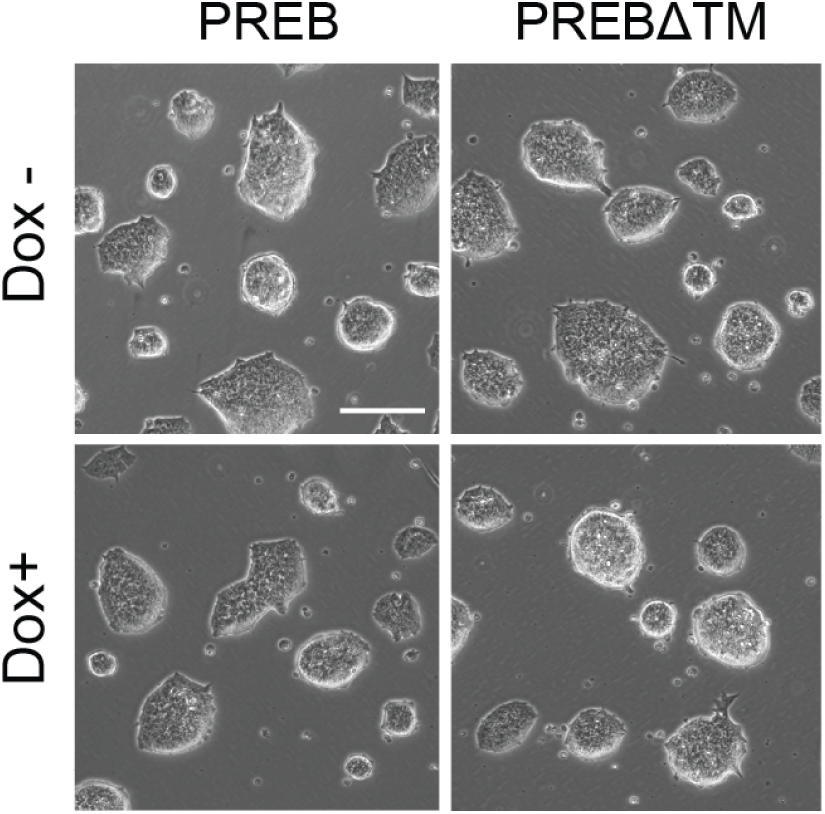
Overexpression of PREB does not change the morphology of mouse ES cells Brightfield images of mouse ES cells expressing full-length PREB and PREBΔTM following treatment with 0.5 µg/mL of Dox for 24 hours (Dox+). Untreated cells (Dox-) were used as controls (Scale bar: 100 µm).

**Supplementary Figure S2.**
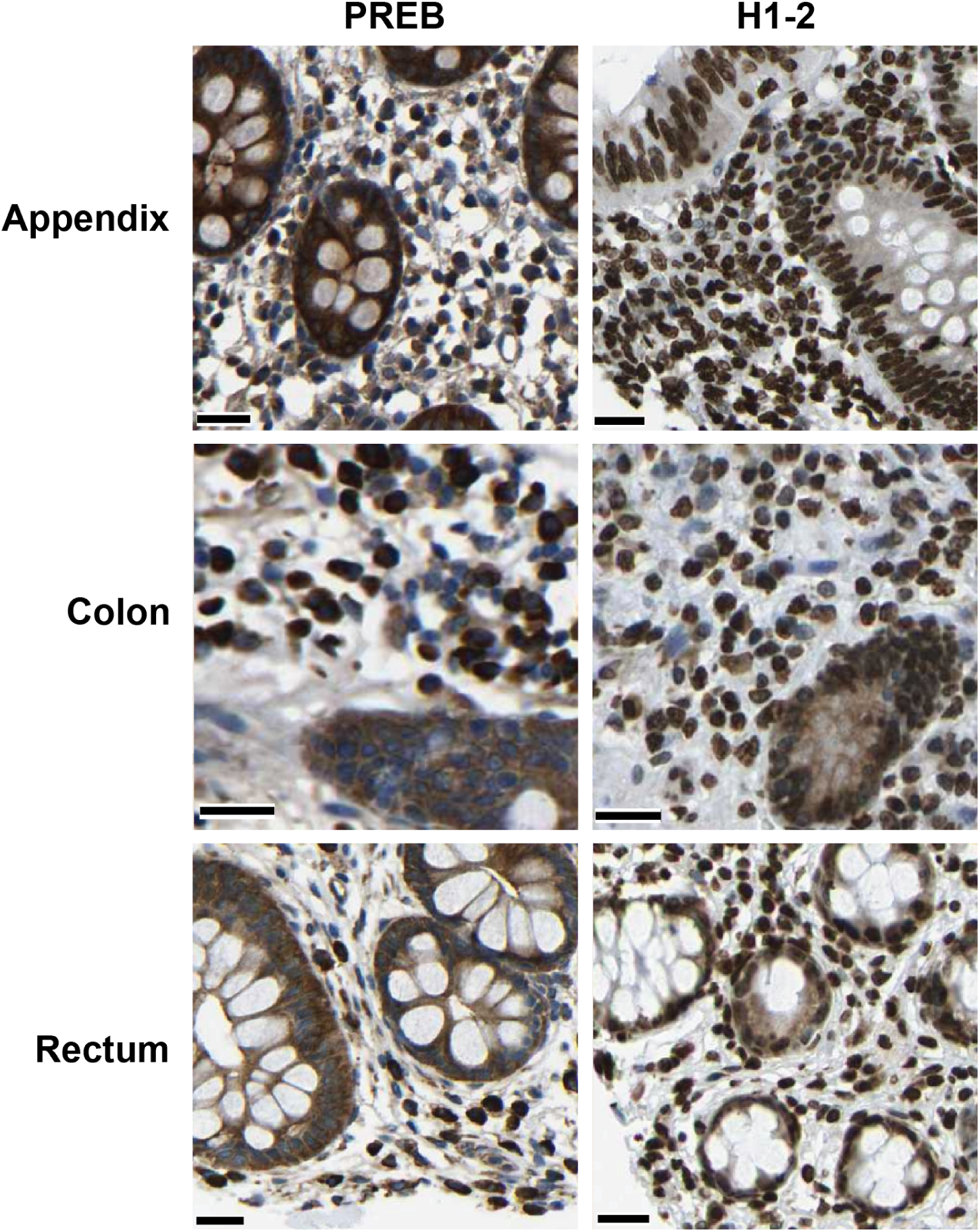
Endogenous PREB localizes to the nucleus in cells of the human appendix, colon, and rectum. Immunohistochemistry-based protein expression profiles of PREB and H1-2 (H1.2 linker histone) in human appendix, colon, and rectum tissues (scale bar = 25 µm), showing data from the Human Protein Atlas (63). The brown 3,3’-diaminobenzidine staining labels target proteins; the blue hematoxylin preferentially labels cell nuclei. These three profiles suggest nuclear localization of PREB in many cells in tissue sections of human appendix, colon, and rectum, but with some variation in localization across specific cell types. In comparison, the histone H1-2 exhibits nuclear localization in almost all cells in tissue-matched sections.

**Supplementary Figure S3.**
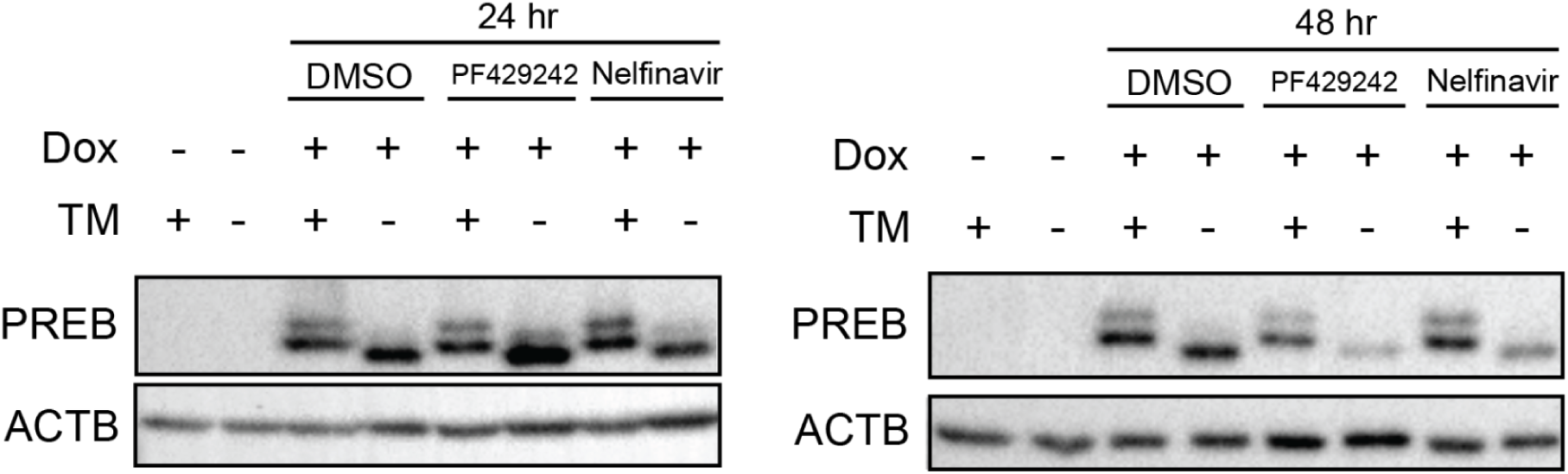
S1P and S2P inhibitors do not block the cleavage of PREB Western blot results after applying S1P and S2P inhibitors for 24 and 48 hours. PREB clones retain two bands after inhibitor treatment, indicating that the two bands do not derive from S1P or S2P protease cleavage activity.

## Supplementary Tables

**Supplementary Table S1.**
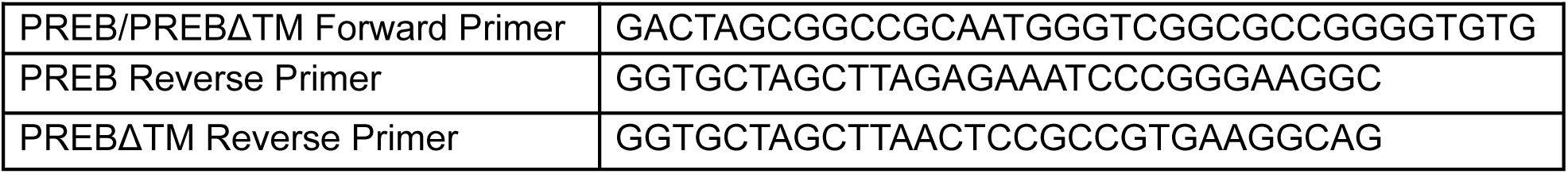
Primers used for cloning.

**Supplementary Table S2. Selected 855 [domain, orthogroup] pairs and corresponding proteins** Supplementary Table 2 is available for download at https://doi.org/10.5281/zenodo.17351618.

## References

1. Latchman, D.S. (1997) Transcription factors: an overview. Int. J. Biochem. Cell Biol., 29, 1305–1312.

2. Dorotea, D., Koya, D. and Ha, H. (2020) Recent Insights Into SREBP as a Direct Mediator of Kidney Fibrosis via Lipid-Independent Pathways. Front. Pharmacol., 11, 265.

3. Li, Z., Park, Y. and Marcotte, E.M. (2013) A Bacteriophage tailspike domain promotes self-cleavage of a human membrane-bound transcription factor, the myelin regulatory factor MYRF. PLoS Biol., 11, e1001624.

4. Glembotski, C.C., Arrieta, A., Blackwood, E.A. and Stauffer, W.T. (2020) ATF6 as a Nodal Regulator of Proteostasis in the Heart. Front. Physiol., 11, 267.

5. Brown, M.S., Ye, J., Rawson, R.B. and Goldstein, J.L. (2000) Regulated Intramembrane Proteolysis. Cell, 100, 391–398.

6. Hoppe, T., Rape, M. and Jentsch, S. (2001) Membrane-bound transcription factors: regulated release by RIP or RUP. Curr. Opin. Cell Biol., 13, 344–348.

7. Zupicich, J., Brenner, S.E. and Skarnes, W.C. (2001) Computational prediction of membrane-tethered transcription factors. Genome Biol., 2, RESEARCH0050.

8. Mistry, J., Chuguransky, S., Williams, L., Qureshi, M., Salazar, G.A., Sonnhammer, E.L.L., Tosatto, S.C.E., Paladin, L., Raj, S., Richardson, L.J., et al. (2021) Pfam: The protein families database in 2021. Nucleic Acids Res., 49, D412–D419.

9. Yao, S., Deng, L. and Zeng, K. (2017) Genome-wide in silico identification of membrane-bound transcription factors in plant species. PeerJ, 5, e4051.

10. Jin, J., Tian, F., Yang, D.-C., Meng, Y.-Q., Kong, L., Luo, J. and Gao, G. (2017) PlantTFDB 4.0: toward a central hub for transcription factors and regulatory interactions in plants. Nucleic Acids Res., 45, D1040–D1045.

11. Itoh, M., Nacher, J.C., Kuma, K., Goto, S. and Kanehisa, M. (2007) Evolutionary history and functional implications of protein domains and their combinations in eukaryotes. Genome Biol., 8, R121.

12. Forslund, K. and Sonnhammer, E.L.L. (2012) Evolution of protein domain architectures. Methods Mol. Biol. Clifton NJ, 856, 187–216.

13. UniProt Consortium (2025) UniProt: the Universal Protein Knowledgebase in 2025. Nucleic Acids Res., 53, D609–D617.

14. Bernhofer, M. and Rost, B. (2022) TMbed: transmembrane proteins predicted through language model embeddings. BMC Bioinformatics, 23, 326.

15. Huerta-Cepas, J., Szklarczyk, D., Heller, D., Hernández-Plaza, A., Forslund, S.K., Cook, H., Mende, D.R., Letunic, I., Rattei, T., Jensen, L.J., et al. (2019) eggNOG 5.0: a hierarchical, functionally and phylogenetically annotated orthology resource based on 5090 organisms and 2502 viruses. Nucleic Acids Res., 47, D309–D314.

16. Cantalapiedra, C.P., Hernández-Plaza, A., Letunic, I., Bork, P. and Huerta-Cepas, J. (2021) eggNOG-mapper v2: Functional Annotation, Orthology Assignments, and Domain Prediction at the Metagenomic Scale. Mol. Biol. Evol., 38, 5825–5829.

17. Katoh, K. and Standley, D.M. (2013) MAFFT Multiple Sequence Alignment Software Version 7: Improvements in Performance and Usability. Mol. Biol. Evol., 30, 772–780.

18. Price, M.N., Dehal, P.S. and Arkin, A.P. (2010) FastTree 2 – Approximately Maximum-Likelihood Trees for Large Alignments. PLoS ONE, 5, e9490.

19. Markin, A., Wagle, S., Grover, S., Vincent Baker, A.L., Eulenstein, O. and Anderson, T.K. (2023) PARNAS: Objectively Selecting the Most Representative Taxa on a Phylogeny. Syst. Biol., 72, 1052–1063.

20. Jones, P., Binns, D., Chang, H.-Y., Fraser, M., Li, W., McAnulla, C., McWilliam, H., Maslen, J., Mitchell, A., Nuka, G., et al. (2014) InterProScan 5: genome-scale protein function classification. Bioinformatics, 30, 1236–1240.

21. Letunic, I. and Bork, P. (2024) Interactive Tree of Life (iTOL) v6: recent updates to the phylogenetic tree display and annotation tool. Nucleic Acids Res., 52, W78–W82.

22. Lee, M., Guo, Q., Kim, M., Choi, J., Segura, A., Genceroglu, A., LeBlanc, L., Ramirez, N., Jang, Y.J., Jang, Y., et al. (2024) Systematic mapping of TF-mediated cell fate changes by a pooled induction coupled with scRNA-seq and multi-omics approaches. Genome Res., 10.1101/gr.277926.123.

23. Kim, J., Cantor, A.B., Orkin, S.H. and Wang, J. (2009) Use of in vivo biotinylation to study protein–protein and protein–DNA interactions in mouse embryonic stem cells. Nat. Protoc., 4, 506–517.

24. Langmead, B. and Salzberg, S.L. (2012) Fast gapped-read alignment with Bowtie 2. Nat. Methods, 9, 357–359.

25. Danecek, P., Bonfield, J.K., Liddle, J., Marshall, J., Ohan, V., Pollard, M.O., Whitwham, A., Keane, T., McCarthy, S.A., Davies, R.M., et al. (2021) Twelve years of SAMtools and BCFtools. GigaScience, 10, giab008.

26. Zhang, Y., Liu, T., Meyer, C.A., Eeckhoute, J., Johnson, D.S., Bernstein, B.E., Nusbaum, C., Myers, R.M., Brown, M., Li, W., et al. (2008) Model-based Analysis of ChIP-Seq (MACS). Genome Biol., 9, R137.

27. Ramírez, F., Ryan, D.P., Grüning, B., Bhardwaj, V., Kilpert, F., Richter, A.S., Heyne, S., Dündar, F. and Manke, T. (2016) deepTools2: a next generation web server for deep-sequencing data analysis. Nucleic Acids Res., 44, W160–165.

28. Jumper, J., Evans, R., Pritzel, A., Green, T., Figurnov, M., Ronneberger, O., Tunyasuvunakool, K., Bates, R., Žídek, A., Potapenko, A., et al. (2021) Highly accurate protein structure prediction with AlphaFold. Nature, 596, 583–589.

29. Amemiya-Kudo, M., Shimano, H., Hasty, A.H., Yahagi, N., Yoshikawa, T., Matsuzaka, T., Okazaki, H., Tamura, Y., Iizuka, Y., Ohashi, K., et al. (2002) Transcriptional activities of nuclear SREBP-1a,-1c, and-2 to different target promoters of lipogenic and cholesterogenic genes. J. Lipid Res., 43, 1220–1235.

30. Blum, M., Andreeva, A., Florentino, L.C., Chuguransky, S.R., Grego, T., Hobbs, E., Pinto, B.L., Orr, A., Paysan-Lafosse, T., Ponamareva, I., et al. (2025) InterPro: the protein sequence classification resource in 2025. Nucleic Acids Res., 53, D444–D456.

31. Yuan, C., Ni, L., Zhang, C. and Wu, X. (2020) The Role of Notch3 Signaling in Kidney Disease. Oxid. Med. Cell. Longev., 2020, 1–8.

32. Hien, L.T. and Back, S.H. (2021) Establishment of a reporter system for monitoring activation of the ER stress transducer ATF6β. Biochem. Biophys. Res. Commun., 558, 1–7.

33. Saito, A., Omura, I. and Imaizumi, K. (2024) CREB3L1 / OASIS: cell cycle regulator and tumor suppressor. FEBS J., 291, 4853–4866.

34. Zhang, K., Shen, X., Wu, J., Sakaki, K., Saunders, T., Rutkowski, D.T., Back, S.H. and Kaufman, R.J. (2006) Endoplasmic Reticulum Stress Activates Cleavage of CREBH to Induce a Systemic Inflammatory Response. Cell, 124, 587–599.

35. Sampieri, L., Di Giusto, P. and Alvarez, C. (2019) CREB3 Transcription Factors: ER-Golgi Stress Transducers as Hubs for Cellular Homeostasis. Front. Cell Dev. Biol., 7, 123.

36. Becker-Herman, S., Arie, G., Medvedovsky, H., Kerem, A. and Shachar, I. (2005) CD74 Is a Member of the Regulated Intramembrane Proteolysis-processed Protein Family. Mol. Biol. Cell, 16, 5061–5069.

37. Gil-Yarom, N., Radomir, L., Sever, L., Kramer, M.P., Lewinsky, H., Bornstein, C., Blecher-Gonen, R., Barnett-Itzhaki, Z., Mirkin, V., Friedlander, G., et al. (2017) CD74 is a novel transcription regulator. Proc. Natl. Acad. Sci. U. S. A., 114, 562–567.

38. Hachmeister, M., Bobowski, K.D., Hogl, S., Dislich, B., Fukumori, A., Eggert, C., Mack, B., Kremling, H., Sarrach, S., Coscia, F., et al. (2013) Regulated Intramembrane Proteolysis and Degradation of Murine Epithelial Cell Adhesion Molecule mEpCAM. PLoS ONE, 8, e71836.

39. May, P., Reddy, Y.K. and Herz, J. (2002) Proteolytic Processing of Low Density Lipoprotein Receptor-related Protein Mediates Regulated Release of Its Intracellular Domain. J. Biol. Chem., 277, 18736–18743.

40. Polavarapu, R., An, J., Zhang, C. and Yepes, M. (2008) Regulated Intramembrane Proteolysis of the Low-Density Lipoprotein Receptor-Related Protein Mediates Ischemic Cell Death. Am. J. Pathol., 172, 1355–1362.

41. Zhang, H., Ma, Q., Zhang, Y.-W. and Xu, H. (2012) Proteolytic processing of Alzheimer’s β-amyloid precursor protein. J. Neurochem., 120 Suppl 1, 9–21.

42. Hébert, S.S., Serneels, L., Tolia, A., Craessaerts, K., Derks, C., Filippov, M.A., Müller, U. and De Strooper, B. (2006) Regulated intramembrane proteolysis of amyloid precursor protein and regulation of expression of putative target genes. EMBO Rep., 7, 739–745.

43. Lammich, S., Okochi, M., Takeda, M., Kaether, C., Capell, A., Zimmer, A.-K., Edbauer, D., Walter, J., Steiner, H. and Haass, C. (2002) Presenilin-dependent Intramembrane Proteolysis of CD44 Leads to the Liberation of Its Intracellular Domain and the Secretion of an Aβ-like Peptide. J. Biol. Chem., 277, 44754–44759.

44. Gires, O., Pan, M., Schinke, H., Canis, M. and Baeuerle, P.A. (2020) Expression and function of epithelial cell adhesion molecule EpCAM: where are we after 40 years? Cancer Metastasis Rev., 39, 969–987.

45. Mao, H., Lockyer, P., Li, L., Ballantyne, C.M., Patterson, C., Xie, L. and Pi, X. (2017) Endothelial LRP1 regulates metabolic responses by acting as a co-activator of PPARγ. Nat. Commun., 8, 14960.

46. Wöhner, B., Li, W., Hey, S., Drobny, A., Werny, L., Becker-Pauly, C., Lucius, R., Zunke, F., Linder, S. and Arnold, P. (2023) Proteolysis of CD44 at the cell surface controls a downstream protease network. Front. Mol. Biosci., 10, 1026810.

47. Guo, A., Wang, Y., Chen, B., Wang, Y., Yuan, J., Zhang, L., Hall, D., Wu, J., Shi, Y., Zhu, Q., et al. (2018) E-C coupling structural protein junctophilin-2 encodes a stress-adaptive transcription regulator. Science, 362, eaan3303.

48. Gao, Y., Nihira, N.T., Bu, X., Chu, C., Zhang, J., Kolodziejczyk, A., Fan, Y., Chan, N.T., Ma, L., Liu, J., et al. (2020) Acetylation-dependent regulation of PD-L1 nuclear translocation dictates the efficacy of anti-PD-1 immunotherapy. Nat. Cell Biol., 22, 1064–1075.

49. Krapivinsky, G., Krapivinsky, L., Manasian, Y. and Clapham, D.E. (2014) The TRPM7 chanzyme is cleaved to release a chromatin-modifying kinase. Cell, 157, 1061–1072.

50. Chubanov, V., Gudermann, T. and Schlingmann, K.P. (2005) Essential role for TRPM6 in epithelial magnesium transport and body magnesium homeostasis. Pflugers Arch., 451, 228–234.

51. Krapivinsky, G., Krapivinsky, L., Renthal, N.E., Santa-Cruz, A., Manasian, Y. and Clapham, D.E. (2017) Histone phosphorylation by TRPM6’s cleaved kinase attenuates adjacent arginine methylation to regulate gene expression. Proc. Natl. Acad. Sci. U. S. A., 114, E7092–E7100.

52. Schapira, M., Tyers, M., Torrent, M. and Arrowsmith, C.H. (2017) WD40 repeat domain proteins: a novel target class? Nat. Rev. Drug Discov., 16, 773–786.

53. Kim, J.L., Mestre, B., Shin, S.-H. and Futerman, A.H. (2021) Ceramide synthases: Reflections on the impact of Dr. Lina M. Obeid. Cell. Signal., 82, 109958.

54. Sociale, M., Wulf, A.-L., Breiden, B., Klee, K., Thielisch, M., Eckardt, F., Sellin, J., Bülow, M.H., Löbbert, S., Weinstock, N., et al. (2018) Ceramide Synthase Schlank Is a Transcriptional Regulator Adapting Gene Expression to Energy Requirements. Cell Rep., 22, 967–978.

55. Karmodiya, K., Anamika, K., Muley, V., Pradhan, S.J., Bhide, Y. and Galande, S. (2014) Camello, a novel family of Histone Acetyltransferases that acetylate histone H4 and is essential for zebrafish development. Sci. Rep., 4, 6076.

56. Fliss, M.S., Hinkle, P.M. and Bancroft, C. (1999) Expression cloning and characterization of PREB (prolactin regulatory element binding), a novel WD motif DNA-binding protein with a capacity to regulate prolactin promoter activity. Mol. Endocrinol. Baltim. Md, 13, 644–657.

57. Ohtsuka, S., Murao, K., Imachi, H., Cao, W.M., Yu, X., Li, J., Iwama, H., Wong, N.C.W., Bancroft, C. and Ishida, T. (2006) Prolactin regulatory element binding protein as a potential transcriptional factor for the insulin gene in response to glucose stimulation. Diabetologia, 49, 1599–1607.

58. Zhang, X.Z., Imachi, H., Lyu, J.Y., Fukunaga, K., Sato, S., Ibata, T., Kobayashi, T., Yoshimoto, T., Kikuchi, F., Dong, T., et al. (2017) Prolactin regulatory element-binding protein is involved in suppression of the adiponectin gene in vivo. J. Endocrinol. Invest., 40, 437–445.

59. Park, J.-M., Kim, M.-Y., Kim, T.-H., Min, D.-K., Yang, G.E. and Ahn, Y.-H. (2018) Prolactin regulatory element-binding (PREB) protein regulates hepatic glucose homeostasis. Biochim. Biophys. Acta Mol. Basis Dis., 1864, 2097–2107.

60. Joiner, A.M.N. and Fromme, J.C. (2021) Structural basis for the initiation of COPII vesicle biogenesis. Struct. Lond. Engl. 1993, 29, 859–872.e6.

61. Ivics, Z., Hackett, P.B., Plasterk, R.H. and Izsvák, Z. (1997) Molecular Reconstruction of Sleeping Beauty, a Tc1-like Transposon from Fish, and Its Transposition in Human Cells. Cell, 91, 501–510.

62. Hackett, P.B., Largaespada, D.A. and Cooper, L.J. (2010) A Transposon and Transposase System for Human Application. Mol. Ther., 18, 674–683.

63. Uhlén, M., Fagerberg, L., Hallström, B.M., Lindskog, C., Oksvold, P., Mardinoglu, A., Sivertsson, Å., Kampf, C., Sjöstedt, E., Asplund, A., et al. (2015) Tissue-based map of the human proteome. Science, 347, 1260419.

64. Danyukova, T., Schöneck, K. and Pohl, S. (2022) Site-1 and site-2 proteases: A team of two in regulated proteolysis. Biochim. Biophys. Acta Mol. Cell Res., 1869, 119138.

65. Wang, T.-B., Geng, M., Jin, H., Tang, A.-G., Sun, H., Zhou, L.-Z., Chen, B.-H., Shen, G. and Sun, Q. (2021) SREBP1 site 1 protease inhibitor PF-429242 suppresses renal cell carcinoma cell growth. Cell Death Dis., 12, 717.

66. Guan, M., Su, L., Yuan, Y.-C., Li, H. and Chow, W.A. (2015) Nelfinavir and nelfinavir analogs block site-2 protease cleavage to inhibit castration-resistant prostate cancer. Sci. Rep., 5, 9698.

67. Lamoureux, J.S., Stuart, D., Tsang, R., Wu, C. and Glover, J.N.M. (2002) Structure of the sporulation-specific transcription factor Ndt80 bound to DNA. EMBO J., 21, 5721–5732.

68. Li, H. and Richardson, W.D. (2016) Evolution of the CNS myelin gene regulatory program. Brain Res., 1641, 111–121.

